# Novel enzymes for amino acid biosynthesis in diverse bacteria and archaea: GapMind 2026

**DOI:** 10.1101/2024.10.14.618325

**Authors:** Morgan N. Price, Valentine V. Trotter, Markus de Raad, Suzanne M. Kosina, Hira P. Lesea, Anthony L. Shiver, Leslie A. Day, Surya Tripathi, Marta Torres, Trenton K. Owens, Audrey Liwen Wang, Anna Lisa Fear, Xuanyu Tao, Jizhong Zhou, Adam M. Deutschbauer, Adam P. Arkin

## Abstract

We previously described GapMind, an automated web-based tool for annotating amino acid biosynthesis pathways in bacterial and archaeal genomes. We used GapMind to identify gaps in biosynthetic pathways and systematically used comparative genomics and high-throughput genetics to identify candidate genes to fill these gaps. We confirmed the activity of ten of the proposed enzymes by using cross-species complementation assays. Highlights include a novel route to glycine, two families that can replace phosphoserine phosphatase, an alternative N-succinyl-L,L-diaminopimelate desuccinylase, an alternative N-acetylornithine deacetylase, and a bifunctional MetB/MetC. We updated GapMind to include these additional enzymes. Across 208 prokaryotes that have high-quality genomes and can grow in minimal media, the average number of unexplained missing steps or gaps in amino acid biosynthesis dropped from 1.4 per genome to 0.7 per genome. The majority of remaining gaps involve the gain or loss of phosphate groups.

## Introduction

Most free-living bacteria can probably make all of the standard amino acids (Ramoneda et al. 2023), but in many of their genomes, the genes for some biosynthetic enzymes cannot be identified. These gaps occur even in the genomes of bacteria and archaea that are experimentally confirmed to be prototrophic (Price et al. 2020). These gaps make it challenging to predict the growth requirements of an organism from its genome sequence, except for well-studied groups such as Enterobacteria or *Pseudomonas* (Seif et al. 2020). For other bacteria, predictions of amino acid auxotrophies are often incorrect (Price et al. 2018a; Price 2023). These gaps in biosynthetic pathways also imply that many novel biosynthetic enzymes remain to be discovered.

We previously described GapMind, a fast web-based tool for annotating amino acid biosynthesis pathways in bacteria and archaea (Price et al. 2020). Because many of the biosynthetic enzymes are not described in standard databases such as Swiss-Prot, MetaCyc, or BRENDA (Caspi et al. 2020; Chang et al. 2021; UniProt Consortium 2023), we curated lists of experimentally-characterized enzymes that can carry out each biosynthetic step. We also included novel enzymes that we identified using randomly-barcoded transposon sequencing (RB-TnSeq) (Price et al. 2018a; Price et al. 2020). Given a genome of interest, GapMind uses these characterized enzymes, along with curated families (hidden Markov models) from TIGRFams (Haft et al. 2013), to identify candidates for each step, at varying levels of confidence. GapMind also uses similarity to proteins with computationally-predicted functions (from Swiss-Prot) to identify additional candidates, but these are never considered high-confidence candidates. Most often, a high-confidence candidate is at least 40% identical to a characterized enzyme, and is less similar to characterized proteins with other functions. GapMind then reports the highest-confidence candidates for each step. Finally, if there are alternate pathways, GapMind selects the pathway that has the most high-confidence candidates and the fewest gaps (steps with only low-confidence candidates or no candidates). An example of GapMind’s results is shown in Figure 1.

**Figure 1:**
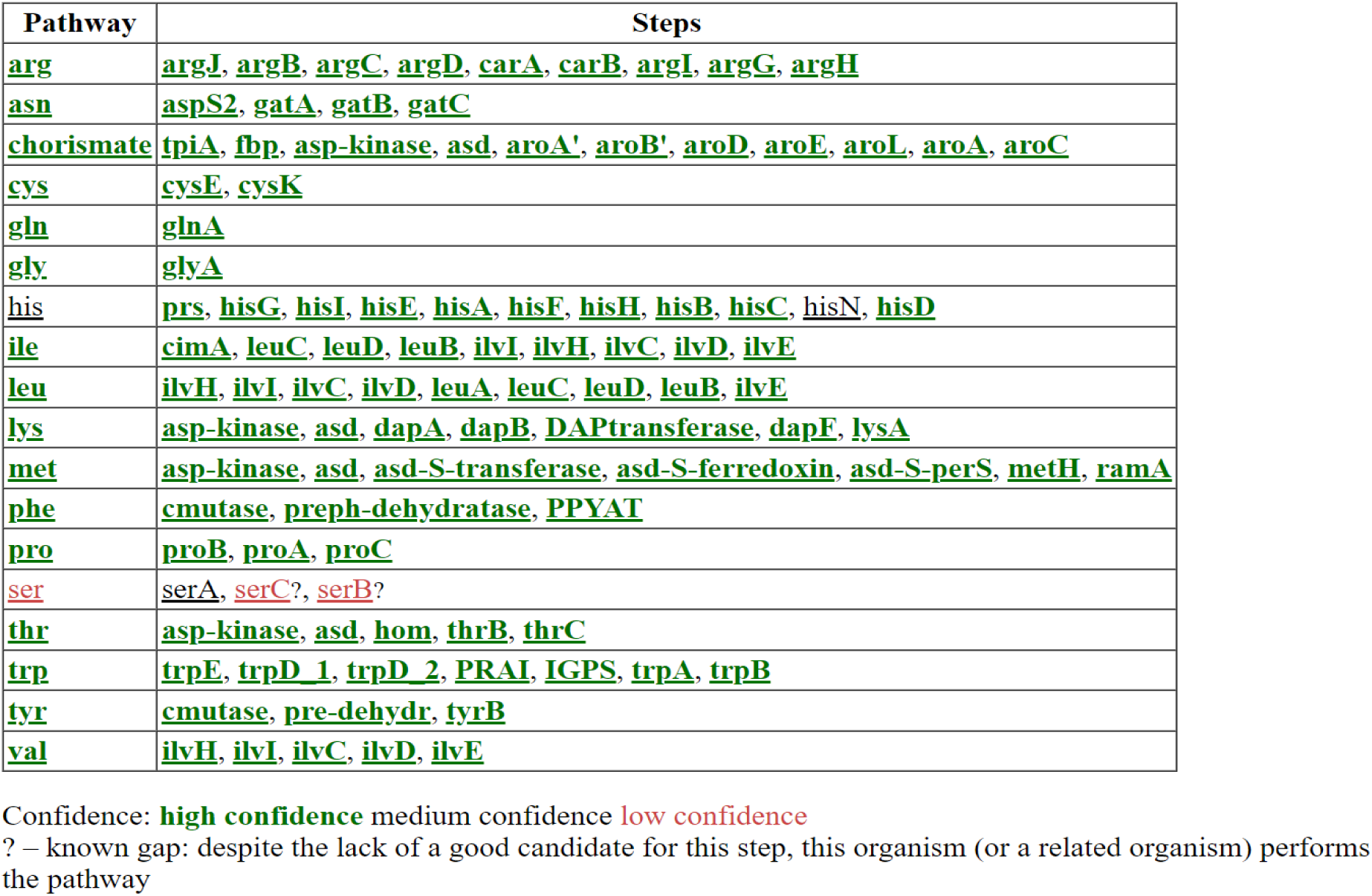
Overview of GapMind’s results for *Nitratidesulfovibrio vulgaris* Hildenborough. In this screenshot, steps and pathways are color-coded by confidence level. All steps are high-confidence except for histidinol-phosphate phosphatase (*hisN*) and the three steps of serine synthesis. Despite these potential gaps, *N. vulgaris* grows in a defined minimal medium (Trotter et al. 2023) and can synthesize all of the standard amino acids.

Since the original publication of GapMind in 2020, over six hundred relevant proteins have been curated in MetaCyc, BRENDA, Swiss-Prot, or the Fitness Browser reannotations (Caspi et al. 2020; Chang et al. 2021; UniProt Consortium 2023; Price and Arkin 2024b). Also, novel enzymes involved in amino acid biosynthesis were predicted based on comparative genomics (Price et al. 2021; Ashniev et al. 2022), and a novel pathway for methionine synthesis was identified in *Streptomyces* (Hasebe et al. 2024). Furthermore, we have collected RB-TnSeq data from many more organisms, including additional phyla of bacteria and archaea (Day et al. 2024; Shiver et al. 2025). As we will show, this allowed us to identify 17 novel or diverged biosynthetic genes. Finally, we used cross-species complementation assays to confirm the functions of 10 other genes. Together, we provide experimental evidence for novel or diverged enzymes involved in the biosynthesis of arginine, chorismate (the precursor to the aromatic amino acids), glycine, isoleucine, leucine, lysine, methionine, serine, and threonine. Based on this new knowledge, we updated GapMind and improved its ability to explain amino acid biosynthesis in diverse bacteria and archaea. GapMind is available at http://papers.genomics.lbl.gov/gaps.

## Results and Discussion

We will first describe the novel genes that we identified, using RB-TnSeq and/or comparative genomics. Second, we will discuss how we incorporated previously-published predictions from comparative genomics into GapMind (Price et al. 2021; Ashniev et al. 2022). Third, we will describe how we improved GapMind’s coverage by annotating diverged enzymes, which would otherwise be identified as low-confidence candidates, or might be missed entirely. Finally, we will assess how well the revised GapMind covers amino acid biosynthesis in prototrophic bacteria and archaea, and, for auxotrophic bacteria, whether GapMind incorrectly identifies complete biosynthetic pathways.

### Novel genes for glycine synthesis

The standard pathway for glycine biosynthesis in bacteria and archaea involves a single enzyme, serine hydroxymethyltransferase (GlyA), which converts serine plus tetrahydrofolate (THF) to glycine plus 5,10-methylene-THF. *Bifidobacterium breve*, a human gut bacterium, and *Methanococcus maripaludis*, a methanogen, are capable of glycine synthesis despite lacking any apparent *glyA* gene. We examined previously-published RB-TnSeq data from both organisms grown in defined media that lack glycine (Day et al. 2024; Shiver et al. 2025). In both organisms, these experiments highlighted two genes which are important for fitness whenever glycine is not available (Figure 2A & 2B). We will call them *glyXL* and *glyXS*. (GlyXL is MMP_RS07345 or BBR_RS12920, which are 61% amino acid identical; GlyXS is MMP_RS03450 or BBR_RS12915, which are 46% identical.) As shown in Figure 2B, in *M. maripaludis*, *glyXL* and *glyXS* are important for fitness in most of the experiments without added glycine, but there are a few exceptions. These exceptions occur because there was relatively little growth in these conditions, and hence relatively little decrease in the relative abundance of auxotrophic mutants. This can be seen by comparing the fitness pattern of *glyXL* to another amino acid biosynthesis gene, *leuA* (Figure 2C). The two patterns are very similar, except that mutants of *glyXL* are rescued by added glycine or (to a lesser extent) by dipeptides that contain glycine.

**Figure 2:**
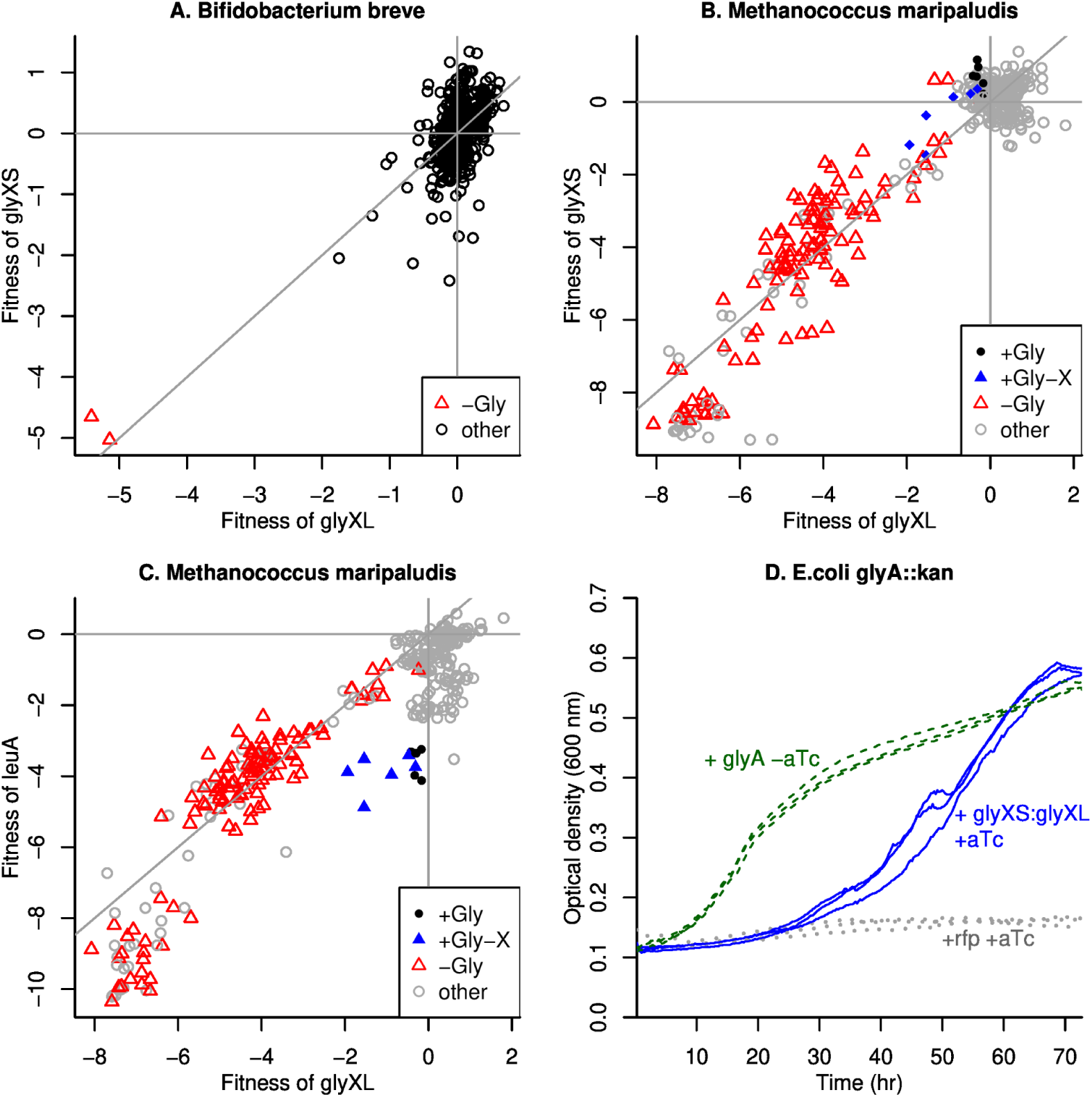
GlyXL and GlyXS are involved in glycine biosynthesis in *Bifidobacterium* and *Methanococcus*. (A,B) We show RB-TnSeq data from *B. breve* UCC2003 (Shiver et al. 2025) and *M. maripaludis* S2 (Day et al. 2024). Each point shows the fitness values of *glyXL* and *glyXS* from a different experiment, where a gene’s fitness is the estimated log2 change in the relative abundance of mutants of that gene during growth (Wetmore et al. 2015). Experiments using a defined medium that lacks glycine are highlighted (red triangles). For *M. maripaludis*, data from defined media with glycine, or with glycine-containing dipeptides (+Gly-X), are also highlighted. (C) Fitness of *glyXL* (*x* axis) and *leuA* (MMP_RS05500, *y* axis) from *M. maripaludis*, with the same color coding as in panel B. In each panel, the lines show *x* = 0, *y* = 0, and *x* = *y*. (D) G*lyA::kan* strains of *E. coli,* carrying various plasmids, were grown in minimal M9 medium with glucose. The *glyA* plasmid carries the gene from *E. coli*; the *glyXS-glyXL* plasmid carries the genes from *B. breve* with the naturally-occurring spacer between them; and the control plasmid carries *rfp* (red fluorescent protein). For strains that express *glyXS-glyXL* or *rfp,* 4 nM of the inducer anhydrotetracycline (aTc) was added.

GlyXL is distantly related to anaerobic ribonucleotide reductases, which use a glycyl radical and two nearby cysteine residues for catalysis. When we aligned the predicted structure of GlyXL from *B. breve* (AlphaFold version 2.0 for UniProt Q8G510; (Varadi et al. 2022)) to the experimental structure of an anaerobic ribonucleotide reductase (PDB:4COM) using the RCSB web site (Bittrich et al. 2024), the structures were not that similar (root mean square deviation 4.38 Å, TM score 0.65), and these catalytic residues were not conserved. So GlyXL may use a different mechanism. GlyXS is an ACT domain protein (PFam PF13740; (Finn et al. 2014)); members of this family often bind amino acids to regulate metabolism.

A role for the GlyXL family in glycine synthesis was previously reported in *Streptococcus pneumoniae* (Kazmierczak et al. 2009). Specifically, *S. pneumoniae* D39 is auxotrophic for glycine and has a truncation in *glyXL* (*spr0218*), and glycine prototrophy can be restored by replacing *glyXL* with the full-length version from a prototrophic strain (Kazmierczak et al. 2009). A complication in interpreting these data is that *S. pneumoniae* D39 also encodes *glyA*. However, metabolite labeling patterns imply that strain D39 uses GlyA in reverse, to convert glycine to serine (Härtel et al. 2012). Furthermore, although strain D39 seems to lack the standard pathway for forming serine (no 3-phosphoglycerate dehydrogenase is apparent in the genome), it does not require serine for growth (Kazmierczak et al. 2009; Härtel et al. 2012); this is consistent with the conversion of glycine to serine.

Across diverse bacteria and archaea, GlyXL and GlyXS usually co-occur and are usually in a putative operon. For example, in the *fast.genomics* database of representative genomes (Price and Arkin 2024a), potential orthologs of GlyXL and GlyXS (above 30% of the best possible bit score) are found in 885 genera, and they are encoded within 5 kb and on the same strand in 815 genera. (But they are not encoded nearby in *M. maripaludis.*) Furthermore, in *Bacillus methanolicus*, expression of the *glyXS-glyXL* operon appears to be regulated by a glycine riboswitch (see BMMGA3_03000 in (Irla et al. 2015)). Although many of the organisms with GlyXL and GlyXS are anaerobic, *B. methanolicus* is obligately aerobic (Arfman et al. 1992), so we predicted that GlyXL-GlyXS would function in the presence of oxygen.

To test if GlyXL-GlyXS can contribute to glycine biosynthesis, we cloned *glyXS* and *glyXL* from *B. breve* into an inducible expression vector (pBbA2c, (Lee et al. 2011)) and introduced them into a *glyA::kan* strain of *E. coli* (Baba et al. 2006). We found that *glyXS-glyXL* enabled growth in minimal M9 medium in the absence of *glyA* (Figure 2D). These experiments were conducted aerobically, which confirms that GlyXL is not a glycyl radical enzyme.

What is the product of GlyXL-GLyXS? Besides these two proteins, fitness data for *B. breve* suggests that glycine synthesis also involves a putative transaminase (BBR_RS14535, fitness = -4.3 or -5.1 in the glycine-free experiments and fitness ≥ -1.1 in the other experiments). So, we propose that BBR_RS14535 is an glyoxylate transaminase (forming glycine) and that GlyXL-GlyXS produce glyoxylate rather than glycine. Although the gene encoding glyoxylate transaminase in *E. coli* is not known, *E. coli* does have an active glyoxylate transaminase (Orsi et al. 2025), so this proposal is consistent with the complementation data. Also, the cofolding prediction tool Boltz-2x (Passaro et al. 2025) predicts that GlyXL from *B. breve* binds glyoxylate much more tightly than it binds glycine (17 μM versus 822 μM). This may be due to a positively-charged binding pocket which includes three arginines (R22, R75, R403) and one negatively-charged residue (E339). We also note that many of the genomes that encode *glyXL* and *glyXS* encode *glyA* as well, but unlike *S. pneumoniae*, most of the genomes with all three of these genes seem to encode the standard pathway for forming serine as well. This is consistent with independent pathways with different starting points for glycine synthesis. In the revised GapMind, glycine can be formed by GlyXL-GlyXS along with a glyoxylate transaminase, and BBR_RS14535 is annotated as a glyoxylate transaminase.

### An alternative to phosphoserine phosphatase from the DUF1015 family

In most prokaryotes, serine is formed from 3-phosphoglycerate, which is an intermediate of glycolysis or gluconeogenesis, in three steps (Figure 3A): 3-phosphoglycerate dehydrogenase (SerA), phosphoserine transaminase (SerC) in reverse, and phosphoserine phosphatase (SerB). No phosphoserine phosphatase (SerB) is apparent in the genome of *Clostridioides difficile* strain 630*Δerm*, but cell extracts of *C. difficile* can convert 3-phosphoglycerate to serine (Hofmann et al. 2018). In *C. difficile*, SerA and SerC are encoded in an operon with a DUF1015 family protein, CDIF630erm_01132 (ARE61905.1), so it was proposed that CDIF630erm_01132 is the missing phosphatase (Hofmann et al. 2018).

**Figure 3:**
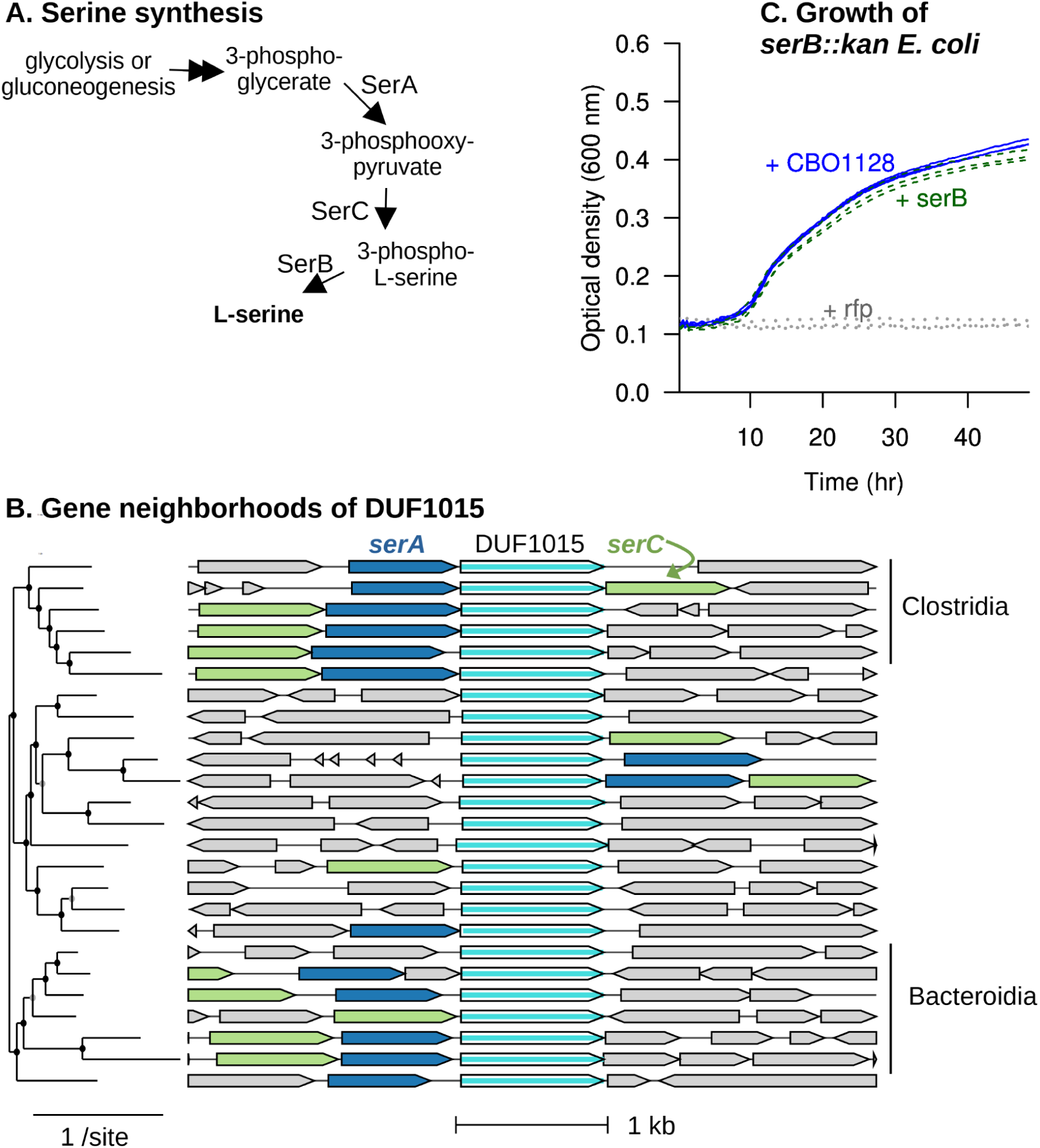
An alternative to phosphoserine phosphatase (SerB) from the DUF1015 family. (A) The standard pathway for serine synthesis. (B) Each row shows a homolog of the putative alternative enzyme (central column) from a different genus, along with its nearby genes. Genes for *serA* and *serC* are color coded. Homologs from *fast.genomics* (Price and Arkin 2024a) were randomly selected from among those whose similarity to CDIF630erm_01132 is at least 30% of the maximum bit score. The phylogenetic tree (from *fast.genomics*) shows the relationships between the DUF1015 proteins. (C) Growth of *serB::kan* strains of *E. coli* carrying various plasmids in minimal M9 medium. (No inducer was added.) CBO1128 is a DUF1015 protein, *serB* was taken from *E. coli*, and *rfp* was used as a control.

Given this hypothesis, we examined the gene neighbors of CDIF630erm_01132 in diverse bacteria. As shown in Figure 3B, homologs of CDIF630erm_01132 are often encoded near *serA*, and are often encoded near a likely *serC* as well. Also, if CDIF630erm_01132 is a replacement for SerB, then its close homologs should be found in genomes that lack the previously-known forms of SerB. We randomly selected 50 representative genomes (all from different genera) that have relatively high-scoring homologs of the putative phosphatase (at least 445 bits, corresponding to roughly 50% identity or above) encoded near *serA*. We ran the updated version of GapMind on these 50 genomes and it identified high-confidence candidates for SerB (at least 40% amino acid identity to a characterized protein) in just three of those 50 genomes. (One of these three was a metagenome-assembled genome, GCA_022072225.1, that also contained two *serA* genes, so it may reflect contamination.) Overall, the vast majority of the genomes that encode close homologs of CDIF630erm_01132 in proximity to *serA* lack *serB*, which again suggests that CDIF630erm_01132 could be a replacement for SerB.

To test this hypothesis experimentally, we cloned the corresponding gene from *Clostridium_F_ botulinum* (CBO1128, whose amino acid sequence is 50% identical to CDIF630erm_01132) into the expression vector pBbA2c. We found that CBO1128 allowed a *serB::kan* strain of *E. coli* (Baba et al. 2006) to grow in minimal medium (Figure 3C).

What is the molecular function of the DUF1015 family? The original study suggested that CDIF630erm_01132 “shows weak homologies to hydrolases” (Hofmann et al. 2018), but we did not identify any sequence similarity to characterized proteins. To identify structural homologs, we used the predicted structure (AlphaFold version 2.0 for UniProt A0A6B4WGC7; (Varadi et al. 2022)) as a query in Foldseek (van Kempen et al. 2024). The top hit from Foldseek was the predicted structure of serine kinase SbnI (UniProt Q2G1M5, root mean square deviation 7.2, e-value 2.7 · 10^-7^). Although this homology is remote, the structural alignment suggested that the ATP binding and catalytic residues are conserved. Also, when we docked CDIF630erm_01132 with ATP using AlphaFold 3 (Abramson et al. 2024), it made a confident prediction of a binding site (interface predicted template modeling score ipTM = 0.91). Thus, even though this family can replace *serB*, they may be kinases rather than phosphatases.

The human gut bacterium *Bacteroides thetaiotaomicron* VPI-5482 encodes a DUF1015 protein, BT1151, which is 47% identical to CDIF630erm_01132. As discussed in Appendix 1, we found that BT1151 was important for the utilization of L-serine as a nitrogen source (gene fitness = -2.2 to -2.6), but it is not important for fitness in most other conditions (data of (Liu et al. 2021)). It is difficult to see how phosphatase activity would support the utilization of serine. In contrast, serine kinase would form phosphoserine, which could be transformed by SerC to 3-phosphohydroxypyruvate and then by SerA (in reverse) to 3-phosphoglycerate, which is an intermediate in glycolysis. A similar pathway for serine utilization was proposed in *Thermococcus kodakarensis* (but not involving DUF1015; (Makino et al. 2016)). Also, although *B. thetaiotaomicron* encodes *serB* (BT0832), *serB* is not important for fitness in most defined media conditions (median fitness = -0.2 across 116 experiments, data of (Liu et al. 2021)). It is also striking that BT1151 is encoded just downstream of *serA* (BT1152) and they are apparently co-transcribed, as no transcript start was identified between BT1151 and BT1152 using Terminator Exonuclease-treated RNA-Seq (Ryan et al. 2024). The simplest explanation is that the BT1151 can replace *serB* for serine synthesis, despite being a kinase rather than a phosphatase.

We propose that CDIF630erm_01132, CBO1128, BT1151, and related proteins from DUF1015 are serine kinases that can participate in both serine synthesis and serine utilization. We named this family *SerBK*, as it seems to provide the capabilities of both the phosphatase (SerB) and the kinase (SerK). However, it is not clear how the serine kinase reaction in reverse could yield sufficient flux to serine for growth, as this is thermodynamically unfavorable (estimated equilibrium constant of 4 ⋅ 10^-4^, (Flamholz et al. 2012)). Furthermore, growing cells usually have a higher concentration of ATP than ADP, which would push the equilibrium further towards phosphoserine. Another uncertainty is that previously-identified serine kinases are either ATP-dependent or ADP-dependent (Makino et al. 2016; Verstraete et al. 2018). Biochemical studies will be necessary to clarify the molecular function of SerBK.

### An alternative phosphoserine phosphatase from the inositol monophosphatase family

In various strains of *Rhodanobacter* and in *Dyella japonica* UNC79MFTsu3.2, which belongs to a related genus, the original GapMind did not identify a convincing candidate for *serB*. However, we noticed that these genomes encode two putative histidinol-phosphate phosphatases (*hisN*). Each genome encodes a *hisN* fused to *hisB* (imidazoleglycerol-phosphate dehydratase), which is presumably annotated correctly, as well as another potential histidinol-phosphate phosphatase from the inositol monophosphatase (IMPase) family. The IMPase family proteins are 38-40% identical to the primary histidinol-phosphate phosphatase (Cg0910) of *Corynebacterium glutamicum* (Kulis-Horn et al. 2017). However, since both *hisBN* and the IMPase family protein are important for fitness in minimal medium (for instance, in *Rhodanobacter denitrificans* FW104-10B01 and in *D. japonica*), we inferred that these IMPase-like proteins have another function. In particular, we predicted that these enzymes would be the missing phosphoserine phosphatases.

To test this hypothesis, we collected RB-TnSeq data for *Rhodanobacter sp.* FW510-R12 in minimal glucose medium, with or without added L-serine. As shown in Figure 4, two of the genes whose fitness was most increased by the addition of L-serine were *serA* (LRK53_RS06795) and the IMPase-like protein (LRK53_RS05785). This confirms that these IMPase-like proteins are alternative phosphoserine phosphatases.

**Figure 4:**
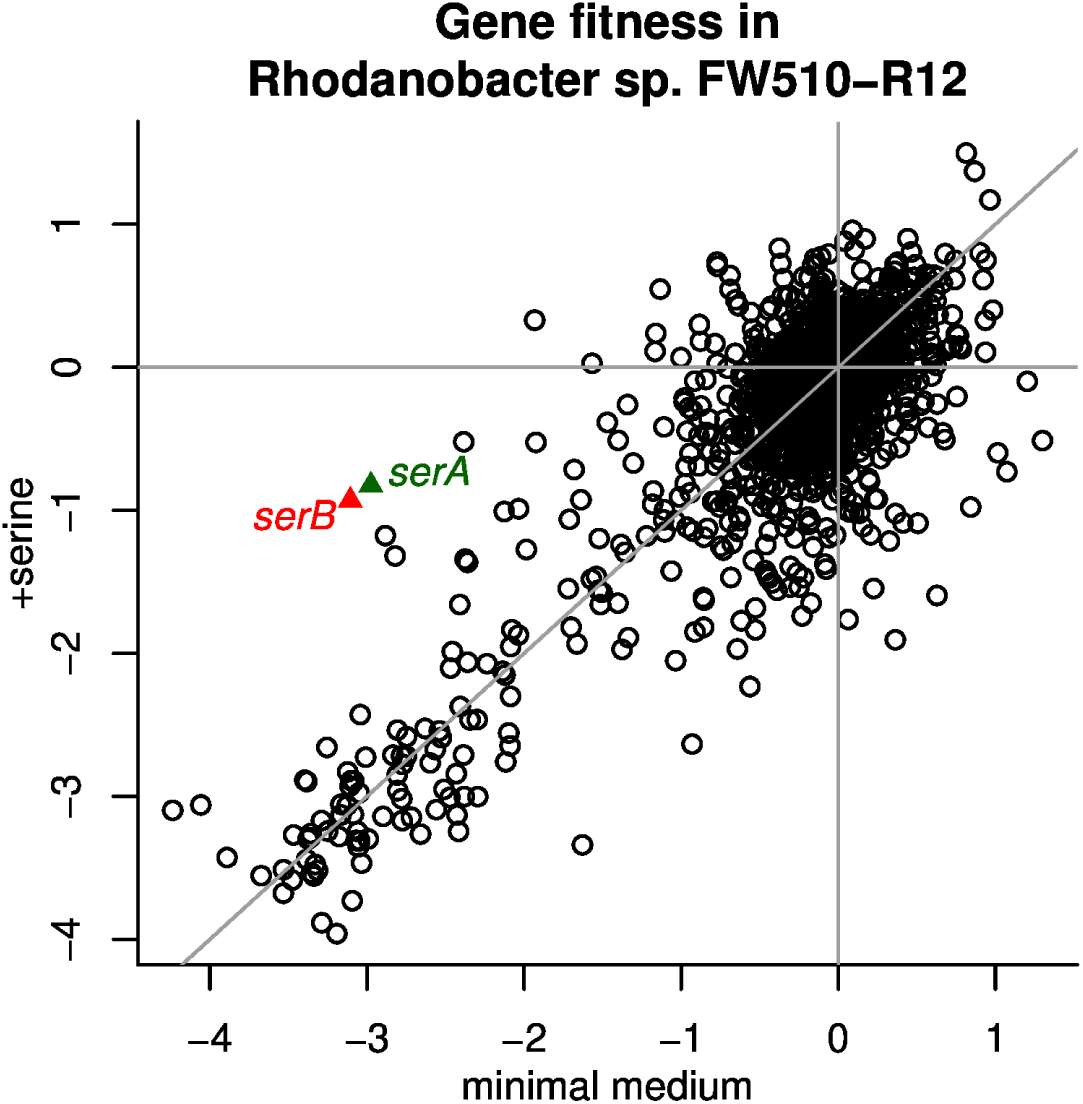
Genetic evidence for an alternative phosphoserine phosphatase (*serB*) in *Rhodanobacter sp.* FW510-R12. Each point shows the average fitness of the gene across two replicate RB-TnSeq experiments, either from minimal glucose medium or from minimal glucose medium with 1 mM L-serine. *SerA* (LRK53_RS06795) and the IMPase-like *serB* (LRK53_RS05785) are highlighted.

### An alternative N-succinyl-L,L-diaminopimelate desuccinylase

In most bacteria, the N-succinyl-L,L-diaminopimelate desuccinylase DapE is required for the synthesis of diaminopimelate (DAP), which is a precursor to both lysine and peptidoglycan. DapE is missing in several prototrophic members of the phylum Bacteroidota that encode the tetrahydrodipicolinate succinylase DapD and hence are expected to use the N-succinyl-DAP pathway (*Echinicola vietnamensis* DSM 17526, *Mucilaginibacter yixingensis* YX-36, and *Pedobacter sp.* GW460-11-11-14-LB5). Because peptidoglycan synthesis is essential for growth, the gene that replaces *dapE* is expected to be essential. Using RB-TnSeq data for all three of these bacteria ((Price et al. 2018b; Torres et al. 2025); see Methods), we searched for conserved essential genes that did not have known functions, and we identified the putative amidohydrolase Echvi_1427 as a candidate to replace DapE (Figure 5A-C). The homolog from another genus of Bacteroidota, *Pontibacter actiniarum* KMM 6156, is also essential (Figure 5D; data of (Price et al. 2018b)).

**Figure 5:**
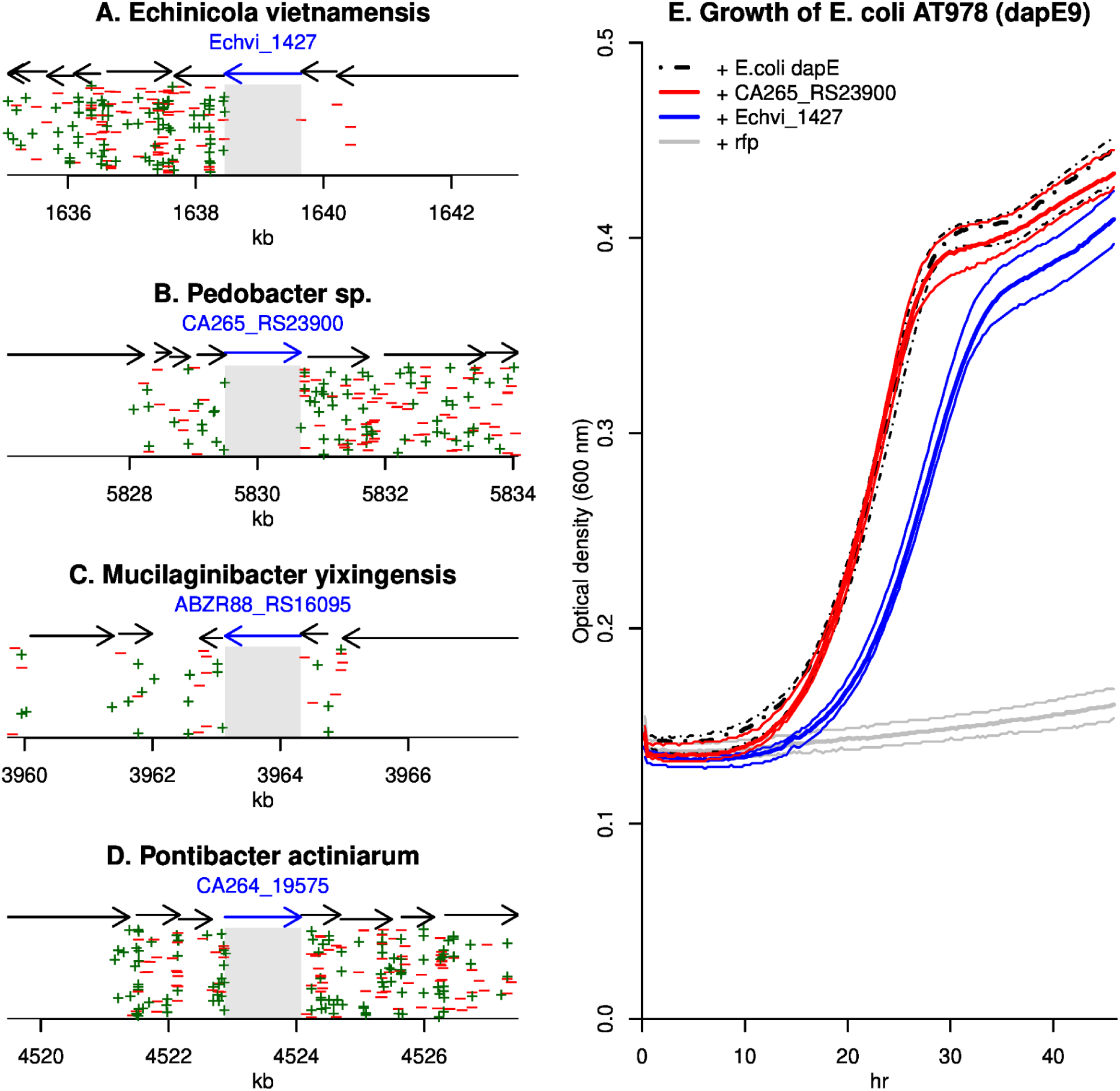
An essential amidohydrolase is an alternative to *dapE*. (A-D) The locations of transposon insertions near Echvi_1427 or its homologs. A “+” symbol indicates that the antibiotic resistance marker was inserted on the forward strand. The region covered by the putative amidohydrolase gene is highlighted in grey, and the *y* axis is random. (E) Growth of *E. coli* AT978, which carries the *dapE9* allele, in M9 minimal medium when carrying various plasmids. Each thick line is the median of six replicates, while thin lines show the minimum and maximum values.

Echvi_1427 is similar to several characterized enzymes that cleave amide bonds, with 42%-43% identity to metal-dependent dipeptidases (Ishikawa et al. 2001; Jamdar et al. 2015) and more distant homology to N-acetyl-L,L-diaminopimelate deacetylase. Echvi_1427 has conserved the metal-binding residues as well as an arginine that binds a terminal carboxylate of the substrate (R264 aligns to R260 from 4EWT; (Jamdar et al. 2015)). These features are consistent with activity on N-succinyl-L,L-diaminopimelate.

To test the role of Echvi_1427 and its homolog CA265_RS23900 (from *Pedobacter sp.* GW460-11-11-14-LB5), we used AT978, a strain of *E. coli* that is auxotrophic for lysine due to the *dapE9* mutation of *dapE* (Bukhari and Taylor 1971). We sequenced *dapE9* and found that codon 78 (out of 375) was mutated to a stop codon (TGG to TGA). In minimal M9 medium, AT978 carrying a control plasmid (expressing RFP) grew slightly or not at all, while AT978 carrying either Echvi_1427, CA265_RS23900, or wild-type *dapE* from *E. coli* grew robustly (Figure 5E). This confirms that Echvi_1427 and its homologs are the missing N-succinyl-L,L-diaminopimelate desuccinylases. Of the representative genomes of Bacteroidota in *fast.genomics* (Price and Arkin 2024a), the majority (391 of 690) encode orthologs of Echvi_1427 and not *E. coli* DapE (BLAST bit score ratios of at least 0.4 for Echvi_1427 and under 0.3 for E. coli DapE).

### Putative bifunctional MetB/MetC in Xanthomonadales

Methionine biosynthesis requires the synthesis of homocysteine, which is usually formed from activated homoserine. In most Gammaprotebacteria, the homoserine is first activated to O-succinylhomoserine by MetA. Then, O-succinylhomoserine is converted to homocysteine, either by direct sulfhydrylation, which is catalyzed by MetZ, or by transsulfuration, in which MetB (cystathionine gamma-synthase) condenses O-succinylhomoserine with cysteine to form cystathionine and MetC (cystathionine beta-lyase) cleaves cystathionine to yield homocysteine (Figure 6A). However, genetic data from Xanthomonadales suggests that methionine synthesis in these bacteria involves a homolog of MetB, but no MetC-like protein.

**Figure 6:**
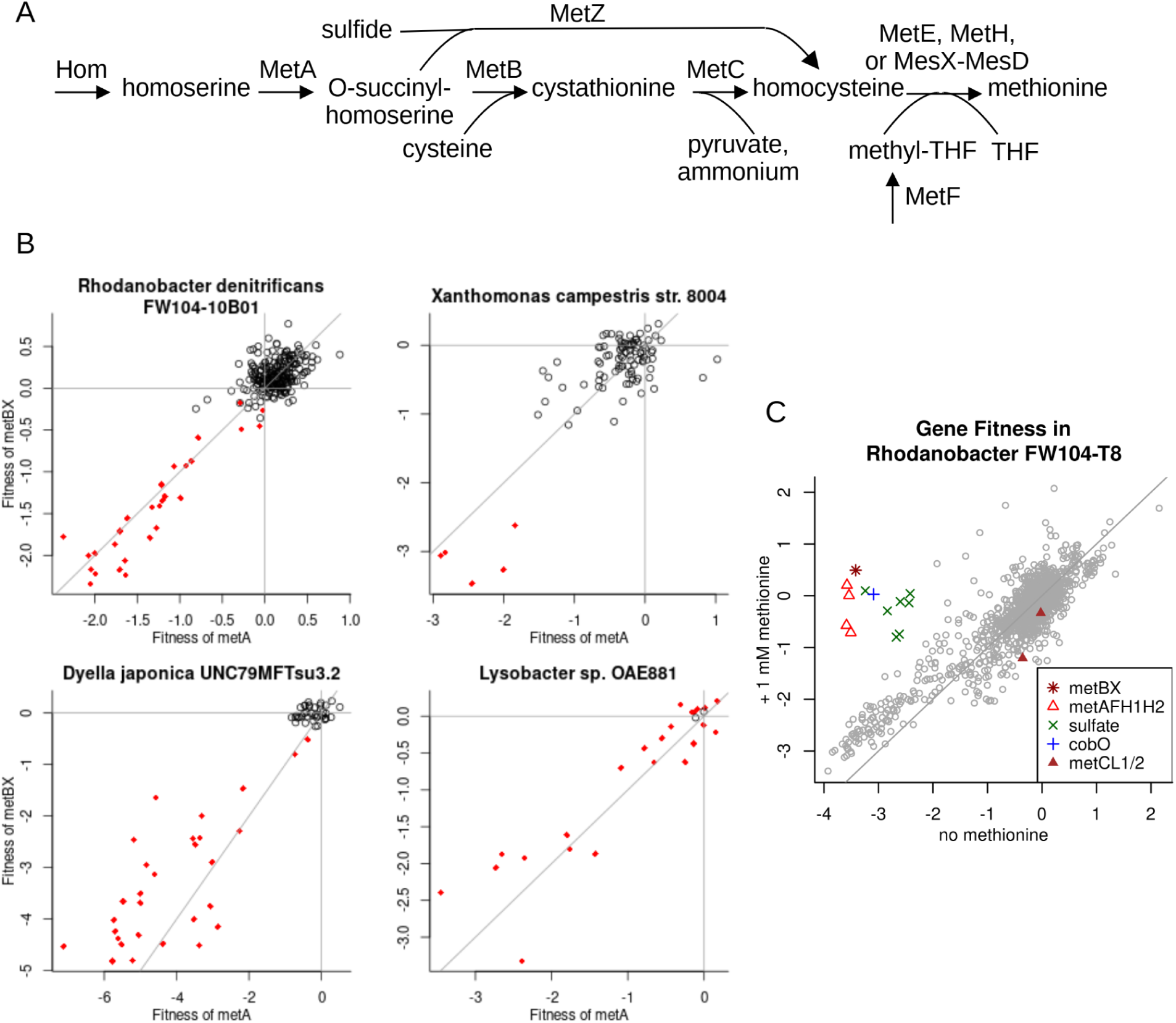
Homocysteine synthesis in Xanthomonadales requires *metBX*. (A) Two alternative pathways for the conversion of O-succinylhomoserine to homocysteine in Gammaproteobacteria. (B) Fitness patterns of *metA* and *metBX* from representatives of four genera of Xanthomonadales. Each point is an independent assay, and experiments conducted in defined media are highlighted in red. For *Rhodanobacter denitrificans* FW104-10B01, the defined medium often included 100 *μ*M of each amino acid (including L-methionine). (C) Fitness data from *Rhodanobacter sp.* FW510-T8 grown in a defined minimal medium with or without 1 mM added L-methionine. The medium included 20 mM D-glucose as a carbon source, ammonium as the nitrogen source, sulfate as the sulfur source, vitamins, and inorganic salts. The *x* axis shows the average of 4 replicate experiments, and the *y* axis shows the average of 3 replicate experiments. Lines show *x* = 0, *y* = 0, or *x = y*. For the identifiers of the genes highlighted in panels B and C, see Supplementary Table S1.

First, in four genera of Xanthomonadales, we identified a *metB*-like gene that has a similar fitness pattern as *metA* (Figure 6B). We named these genes *metBX*, for Xanthomonadales *metB*. (The encoded proteins are 57-60% identical to MetB from *E. coli*.) To quantify the similarity of fitness patterns of *metA* and *metBX*, we use the “cofitness” or linear (Pearson) correlation, which ranged from 0.79 to 0.94. *MetBX* is usually encoded between *metA* and *hom* (which is responsible for the synthesis of homoserine).

Second, we did not find a *metC*-like gene that was involved in methionine synthesis in these four Xanthomonadales. Each of these genomes does encode two MetC-like proteins, which we will call MetCL1 and MetCL2 (for MetC-like). However, none of these genes were cofit with *metA* (cofitness = -0.65 to -0.30) or with *metBX* (cofitness = -0.78 to -0.26). Prior studies confirm that MetCL1 is not involved in methionine synthesis. In *Xanthomonas oryzae,* a mutant of MetCL1 (PXO_03157) is not required for growth in minimal media, and the mutant has a slight deficiency in growth in the absence of cysteine, not methionine (Li et al. 2019). A study of another strain of *X. oryzae* found that a mutant of MetCL1 (XOO0778) had reduced exopolysaccharide and a loss of O antigen (Wang et al. 2008). In a large-scale screen for genes that can complement Δ*metB* or Δ*metC* strains of *E. coli* when overexpressed, the *metCL1* gene from *Rhodanobacter denitrificans* (LRK54_RS05660) did not complement Δ*metC*, although it did complement *ΔmetB* (Biggs et al. 2024). A biochemical study of MetCL1 from *X. oryzae* found that when incubated with cystathionine, it catalyzed the formation of both cysteine and homocysteine (Ngo et al. 2008); if this enzyme were adapted to a role in methionine synthesis, then it should form homocysteine only, as is thought to be the case for MetC from *E. coli* (Farsi et al. 2009). Although we did not find prior data on MetCL2, MetCL2 is often encoded in the same apparent operon as MetCL1, and they have similar fitness patterns in three of the bacteria we studied (cofitness = 0.75 to 0.84; fitness data is not available for *metCL1* in *Dyella japonica*). Overall, it seems that MetCL1 and MetCL2 are involved in cell wall synthesis and are unlikely to provide the missing cystathionine beta-lyase activity.

Third, to search for additional genes that might be involved in methionine synthesis, we assayed a mutant library of *Rhodanobacter sp.* FW510-T8 in minimal media with or without 1 mM of added methionine (Figure 6C). We identified strong effects of methionine on gene fitness for *metA, metBX,* cobalamin-dependent methionine synthase *metH* (split into two genes), and *metF* (which forms 5-methyltetrahydrofolate, the methyl donor for methionine synthase). We also found strong differences for genes that are involved in sulfate assimilation (the enzymes *cysDGHIJNC* and a putative *cysB*-like regulator OKGIIK_05975); sulfate assimilation may no longer be necessary if sulfur is obtained from methionine. The only remaining gene with a strong effect of methionine on its fitness was a putative ATP:cob(I)alamin adenosyltransferase (*cobO*, OKGIIK_11600), which is not explained. We did not find any new candidate genes for methionine synthesis.

Based on the genetic data, it appears that MetBX alone, without the aid of another protein, catalyzes the conversion of O-succinylhomoserine to homocysteine in Xanthomonadales. Because MetB and MetC are related enzymes, MetBX could be bifunctional, with cysteine as the sulfur source and cystathionine as an intermediate. Alternatively, it could be an O-succinylhomoserine sulfhydrylase like MetZ, since MetZ is also similar to MetBX (for instance, MetZ from *Pseudomonas aeruginosa* is 42% identical to MetBX from FW510-T8). Unpublished structures of MetBX from *X. oryzae* in complexes with aminoacrylate and cysteine or with cystathionine (PDB:6LD8, PDB:6LD9) suggest that MetBX could use cysteine as a substrate and that cystathionine could be an intermediate. So, we predict that Xanthomonadales use a single bifunctional protein to perform both steps in the transsulfuration pathway. This would represent the ancestral state before the divergence of MetB and MetC.

### An alternative N-acetylornithine deacetylase

A recent study proposed that an uncharacterized protein in *Veillonella denticariosi* AS16 (UniProt W3Y6L2) is an alternate N-acetylglutamate synthase because of its conserved proximity to arginine synthesis genes (“ArgA3” in (Ashniev et al. 2022)). However, W3Y6L2 is 42% identical to an N-acetyl-cysteine deacetylase (SndA from *Bacillus subtilis*; (Chan et al. 2014; Hazra et al. 2022)), and the only analogous reaction in arginine synthesis is N-acetylornithine deacetylase (ArgE). Indeed, using *fast.genomics*, we found that representative genomes with closer homologs of W3Y6L2 (11 genera contain a homolog with a bit score ratio above 0.4) never contained likely ArgE proteins (homologs of *E. coli* ArgE with a bit score ratio above 0.2). So, we propose that W3Y6L2 is an alternative N-acetylornithine deacetylase.

We cloned W3Y6L2 into an *argE::kan* strain of *E. coli* and found that it rescued growth in minimal medium, but only under anaerobic conditions. During anaerobic growth in M9 medium with glucose and Wolfe’s minerals, the OD_600_ increased by 0.18-0.20 (3 replicates). We did not expect that this enzyme would be sensitive to oxygen, as this family does not use any redox cofactors. Since *V. denticariosi* is strictly anaerobic (Byun et al. 2007), its enzymes might not have evolved to tolerate oxygen. Alternatively, homologs of W3Y6L2 are variously reported to use a Zn^2+^ or Co^2+^ cofactor (SndA, 42% identity, (Hazra et al. 2022)) or a Ni^2+^ cofactor (PDB:1YSJ, 38% identity). Since *E. coli* only uses nickel under anaerobic conditions (for [NiFe] hydrogenases), it is possible that W3Y6L2 requires a Ni^2+^ cofactor which is not available in aerobically-grown *E. coli*.

### A predicted alternative N-succinyl-L-glutamate synthase

In *Steroidobacter denitrificans* DSM 18526, the N-succinyl-L-glutamate synthase (ArgA) is missing. The gene cluster for arginine biosynthesis in *Steroidobacter* and related bacteria includes a very distant homolog (ACG33_RS14135) of a D-glutamate acetyltransferase (Yu et al. 2023), so we predict that ACG33_RS14135 encodes the missing N-succinyl-L-glutamate synthase. For details, see Appendix 2.

### Previously-proposed alternate enzymes

The revised GapMind also includes a number of previously-proposed alternate enzymes (Table 1). Six of these already had experimental evidence for their role; we obtained experimental evidence for three additional enzymes; and the other four await experimental tests. All of the predictions are described in Appendix 3 and Appendix 4; here we briefly describe the new experimental evidence.

**Table 1:** Alternate enzymes that were previously proposed and are included in the revised GapMind.

| Pathway | Step | Evidence | Genus | UniProt | Reference |
| --- | --- | --- | --- | --- | --- |
| Met | Sulfur carrier protein (MetO) | yes | *Streptomyces* | A0A059W7F4 | (Hasebe et al. 2024) |
| Met | [MetO]-L-cysteine hydrolase (Mec) | yes | *Streptomyces* | A0A059W3G7 | (Hasebe et al. 2024) |
| Met | [MetO]-sulfur ligase (MoeZ) | yes | *Streptomyces* | A0A401R754 | (Hasebe et al. 2024) |
| Met | MetO-thiocarboxylate-dependent homocysteine synthase (MetM) | yes | *Streptomyces* | A0A059WBP2 | (Hasebe et al. 2024) |
| Pro | Ornithine cyclodeaminase | yes | *Methanococcus* | Q6LXX7 | (Burnat et al. 2019) |
| Ser | Phosphoserine transaminase (SerC2) | yes | *Staphylococcus* | Q2FXK2 | (Verstraete et al. 2018; Ashniev et al. 2022) |
| Thr | Homoserine phosphotransferase (HomK) | this work | *Bacteroides* | Q8A542 | (Price et al. 2020) |
| Ser | Phosphoserine transaminase (SerC3) | this work | *Clostridium*<sub>F</sub> | A5I0W7 | (Ashniev et al. 2022); this work |
| Lys | Diaminopimelate epimerase (DapF2) | this work | *Staphylococcus* | A0A9Q5JK70 | (Ashniev et al. 2022) |
| Lys | N-acetyl-diaminopimelate deacetylase (DapL) | predicted | *Staphylococcus* | Q2FH40 | (Jiang et al. 2015; Ashniev et al. 2022) |
| Met | Split MetE, folate-binding component | predicted | *Pyrolobus* | G0EDA1 | (Price et al. 2021) |
| Met | Split MetE, catalytic component | predicted | *Pyrolobus* | G0EFB7 | (Price et al. 2021) |
| Met | Corrinoid-dependent methionine synthase (MesC) | predicted | *Methanosarcina* | Q8TUL3 | (Price et al. 2021) |

First, we previously reported that a putative alternative to homoserine kinase (BT2402) is required for threonine synthesis in *Bacteroides thetaiotaomicron* (Price et al. 2020). Close homologs of BT2402 (over 80% identical) are involved in threonine synthesis in the genus *Phocaiecola* (S. Tripathi and A. M. Deutschbauer, in preparation; data of (Huang et al. 2024); Appendix 3). We cloned BT2402 into a thrB::kan strain of *E. coli* and found that it could rescue growth in minimal medium glucose medium with aTc: OD_600_ increased by 0.14 after 47 hours (all three replicates). This confirms that BT2402 and related proteins (also known as TIGR02535 (Haft et al. 2013) or ThrB2 (Ashniev et al. 2022)) are alternatives to homoserine kinase. BT2402 is related to cofactor-independent phosphoglycerate mutases and probably uses a different phosphate donor than ATP, perhaps phosphoserine (Appendix 3). Because we predict that BT2402 catalyzes a different reaction than homoserine kinase (ThrB), it is included in GapMind as a different step. We chose the name HomK, as it is an alternative to the usual homoserine kinase (ThrB).

Second, Ashniev and colleagues associated a family of putative transaminases, which they named SerC3, with serine synthesis (Ashniev et al. 2022). We cloned the representative of this family from *Clostridium_F_ botulinum* (UniProt A5I0W7) into a *serC::kan* strain of *E. coli* (Baba et al. 2006) and found that it rescued growth in minimal medium: in M9 glucose medium with aTc, the OD_600_ increased by 0.16-0.18 after 48 hours (3 replicates). This confirms that *serC3* proteins are phosphoserine transaminases.

Third, Ashniev and colleagues associated a family of putative epimerases, which they named DapF2, with lysine synthesis, and predicted that they are diaminopimelate epimerases (Ashniev et al. 2022). This family includes the Alr2 protein from *Staphylococcus aureus*, which is often annotated as alanine racemase, but Alr2 is not involved in D-alanine synthesis in *S. aureus* (Panda et al. 2024). To test the role of *dapF2*, we cloned the gene from *S. epidermis* (UniProt A0A9Q5JK70 = HMPREF2922_03580) into a Δ*dapF* strain of *E. coli* (see Methods). The Δ*dapF* strain does not have a growth defect, but strains of *E. coli* that lack *dapF* are reported to secrete L,L-diaminopimelate (which is the substrate of DapF) into the medium (Mengin-Lecreulx et al. 1988). It is thought that in Δ*dapF* cells, L,L-diaminopimelate is overproduced, and some other racemase converts L,L-diaminopimelate to *meso*-diaminopimelate (the precursor of both peptidoglycan and lysine) with low efficiency, while the excess L,L-diaminopimelate is secreted (Mengin-Lecreulx et al. 1988). We used mass spectrometry to compare the supernatant from various strains of *E. coli* and confirmed that cells that lack *dapF* secreted far more diaminopimelate than did wild-type cells. Furthermore, expressing either *dapF* from *E. coli* or *dapF2* from *S. epidermis* in the mutant background prevents this secretion (Appendix 4). This confirms that DapF2 is a diaminopimelate epimerase.

### Diverged enzymes

GapMind only considers a protein to be a high-confidence candidate for a step if it is at least 40% identical to a characterized enzyme. (For enzymes that have curated models in TIGRFam, high-confidence candidates can also be identified via the hidden Markov model, but 38% of steps in GapMind are not associated with any TIGRFam, and non-canonical enzymes are usually not described in TIGRFam.) To improve the coverage of GapMind, we searched for diverged enzymes that had experimental evidence but were missing from the curated databases that GapMind relies on. We also used fitness data to confirm seven enzymes, and we used complementation assays to confirm another three (Appendix 5). One of the enzymes that we confirmed by complementation is a phosphoserine transaminase (SerC, UniParc A0A843E9R6) from a metagenome-assembled genome of a Methanomethylophilaceae. As far as we know, this family of archaea has not yet been cultured yet, but we were still able to fill this gap in serine biosynthesis. Overall, we added 19 diverged enzymes with experimental evidence to GapMind (Table 2).

**Table 2:** Diverged enzymes with experimental evidence that are included in the revised GapMind.

| Pathway | Step | Genus | Identifier | Reference |
| --- | --- | --- | --- | --- |
| Arg, Lys | LysZ | *Thermococcus* | TK0276 | (Yoshida et al. 2016) |
| chorismate | AroA | *Bacteroides* | BT_RS11065 | (Biggs et al. 2024) |
| chorismate | AroE | *Sphingomonas* | BDW16_RS10815 | (Biggs et al. 2024) |
| His | HisC | *Acidovorax* | AAFF19_05770 | (Biggs et al. 2024) |
| His | HisN | *Bifidobacterium* | A0A0L7BRC5 | (Shiver et al. 2025) |
| Ile, Val | IlvH | *Rhodanobacter* | LRK54_RS10305 | This work |
| Ile, Val | IlvH | *Lysobacter* | ABIE51_RS17555 | This work |
| Ile, Val | IlvH | *Xanthomonas* | Xcc-8004.1058.1 | This work |
| Met, Thr | Hom | *Brevundimonas* | A4249_RS00005 | This work |
| Phe | Prephenate dehydratase | *Brevundimonas* | A4249_RS06835 | This work |
| Pro | ProB | *Pedobacter* | AAFF35_11135 | (Biggs et al. 2024) |
| Ser | SerB | *Lysobacter* | GGC55_RS00335 | This work |
| Ser | SerB | *Xanthomonas* | A0A0H2XA29 | This work |
| Ser | SerC | *Paucidesulfobivrio* | A0A1T4W7T3 | This work |
| Ser | SerC | *Methanomethylophilaceae* | A0A843E9R6 | This work |
| Thr | ThrB | *Bacillus* | BSU_32240 | (Parsot 1986; Biggs et al. 2024) |
| Thr | HomK | *Dehalococcoides* | DET_RS08350 | This work |
| Trp | TrpA | *Pedobacter* | AAFF35_06155 | (Biggs et al. 2024) |
| Trp | TrpA | *Rhodanobacter* | LRK54_RS01680 | (Biggs et al. 2024) |

We identified another 28 diverged enzymes that lack experimental evidence but are confirmed by conserved gene context and are required to explain the biosynthesis of the amino acid in a prototrophic prokaryote (Table 3). For these predicted enzymes, where possible, we also checked that catalytic residues were present, using SitesBLAST ((Price and Arkin 2022); for functional residues, see Supplementary Table S2). The highly-diverged regulatory subunit for acetolactate synthase (IlvH) from *Korarcheum cryptofilum* was checked using structural analyses (Appendix 6). Three diverged HomK enzymes were identified by the combination of being encoded near *hom* (Table 3) and functional residues (see Appendix 3).

**Table 3:** Diverged enzymes that are included as predictions in the revised GapMind.

| Pathway | Step | Organism | Identifier |
| --- | --- | --- | --- |
| Arg | LysK | *Nitrosopumilus maritimus* | NMAR_RS06940 |
| chorismate | AroD | *Pyrolobus fumarii* | PYRFU_RS04235 |
| chorismate | AroA | *Pyrolobus fumarii* | PYRFU_RS00635 |
| Gln | GlnA | *Clostridium kluyveri* | CKL_RS00630 |
| Lys | LysA | *Chitinovibrio alkaliphilus* | CALK_RS03395 |
| Lys | LysZ | *Haloferax volcanii* | D4GYN9 |
| Ile, Val | IlvH | *Brevundimonas* sp. | A4249_RS10015 |
| Ile, Val | IlvH | *Chryseobacterium* sp. | A0A316WEF5 |
| Ile, Val | IlvH | *Korarchaeum cryptofilum* | KCR_RS03285 |
| Met | Split MetE, folate-binding | *Haloferax volcanii* | D4GW95 |
| Met | Split MetE, catalytic | *Haloferax volcanii* | D4GW90 |
| Met | B12 reactivation | *Heliobacterium modesticaldum* | H1S01_RS06050 |
| Met | L-aspartate semialdehyde<br>S-transferase, NIL/ferredoxin | *Methanococcus vannielii* | MEVAN_RS03425 |
| Met | L-aspartate semialdehyde<br>S-transferase, persulfide-form. | *Hippea alviniae* | G415_RS0107280 |
| Met | Mec | *Dissulfurispira thermophila* | A0A7G1H0A1 |
| Phe | Prephenate dehydratase | *Carboxydotherrmus pertinax* | A0A1L8CUL6 |
| Ser | SerA | *Ideonella sp.* | A0A257CND8 |
| Ser | SerA | *Roseburia faecis* | A0A374I5P9 |
| Ser | SerA | *Rhodoferrax sp.* | A0A975E934 |
| Ser | SerBK | *Sediminibacterium sp.* | A0A521P7R6 |
| Ser | SerC | *Thermoanaerobacter kivui* | A0A097AUI2 |
| Thr | HomK | *Paramuribaculum intestinale* | A0A2V1IX31 |
| Thr | HomK | *Microtrichaceae sp.* | DCC48_11680 |
| Thr | HomK | *Megalodesulfobivrio gigas* | T2GDP6 |
| Tyr | Prephenate dehydrogenase | *Dehalococcoides mccartyi* | DET_RS02500 |
| Tyr | Prephenate dehydrogenase | *Desulfarculus baarsii* | E1QE65 |
| Tyr | Prephenate dehydrogenase | *Desulfobulbus oligotrophicus* | A0A7T5VEL0 |
| Tyr | Prephenate dehydrogenase | *Nitrosopumilus maritimus* | A9A228 |

### The accuracy of GapMind for prototrophs

To test the coverage of the revised GapMind, we analyzed the predicted proteomes of 208 diverse bacteria and archaea that can grow in defined minimal medium (see Methods). We ensured that the growth data and the genome sequence were from the same strain. These 208 prototrophs cover 20 phyla and 163 genera from the Genome Taxonomy Database (Chaumeil et al. 2019). Across 3,744 pathway x organism combinations, 84% were fully complete and consisted only of high-confidence steps. Another 11% of pathways contained one or more medium-confidence steps, such as a comparative genomics prediction or a diverged enzyme. (Unless the step is described by a hidden Markov model, high-confidence candidates must be at least 40% identical to a characterized enzyme.) 5% of pathways had one or more gaps, with no high- or medium-confidence candidate for at least one step.

Overall, we identified 187 gaps in these 208 genomes. However, for 37 of these gaps, a high- or medium-confidence candidate was identified when analyzing the six-frame translation of the genome. Most of these discrepancies were due to missing gene calls, but these cases also include seven distinct frame shifts that led to 13 gaps. (Some steps occur in more than one pathway.) We previously showed that three putative frame shifts in amino acid biosynthesis genes from prototrophic bacteria were actually errors in their genome sequences (Price et al. 2018a; Price et al. 2020). That includes one of the seven frame shifts in our current analysis, but we expect that the other six are spurious as well. This left 150 genuine gaps, corresponding to 0.7% of all steps on the best paths. For example, one of the genomes with the most gaps was *Desulfotalea psychrophila* LSv54, which is a sulfate-reducing bacterium that grows with lactate as the sole carbon source (Rabus et al. 2004). The genome has apparent frameshifts in *ilvC, ilvD*, *aroB*, and L,L-diaminopimelate aminotransferase. The genome was sequenced at only 6.4x coverage (Rabus et al. 2004), which suggests that these frameshifts could be spurious. This genome also has three genuine gaps: phosphoribosyl-ATP pyrophosphatase *(hisI*), histidinol-phosphate phosphatase (*hisN*), and phosphoserine phosphatase (*serB*) are all missing.

Across all 208 prototrophic genomes, if we consider each step once, even if it occurs in multiple pathways, then we have 143 genuine gaps, or an average of 0.7 per genome. In contrast, the original GapMind had 282 genuine gaps for these genomes (an average of 1.4). Just four steps account for 65% of the remaining gaps: histidinol-phosphate phosphatase *hisN* is missing in 42 genomes (20% of prototrophs), phosphoribosyl-ATP pyrophosphatase *hisI* is missing in 19 genomes (9%), phosphoserine phosphatase *serB* is missing in 16 genomes (8%), and homoserine kinase *thrB* is missing in 16 genomes (8%). All four of these enzymes catalyze the gain or loss of phosphate or pyrophosphate groups, which can be carried out by many different protein families. We speculate that other kinases, phosphatases, or pyrophosphatases have evolved to act on these substrates and sometimes replace the known families. In any case, the absence of these genes should not be used to predict auxotrophies.

As shown in Figure 7, most prototrophic genomes from the phylum Pseudomonadota do not have any genuine gaps. This is probably because Pseudomonadota is the best-studied phylum. (Of the 163 known prototrophic genera in our collection, 77 are Pseudomonadota.) In contrast, most of the other bacteria and archaea that we studied have at least one genuine gap in their amino acid biosynthesis pathways. Even for the Pseudomonadota, the true rate of gaps may be higher, as we used the same set of prototrophic genomes to help us fill gaps in biosynthetic pathways.

**Figure 7:**
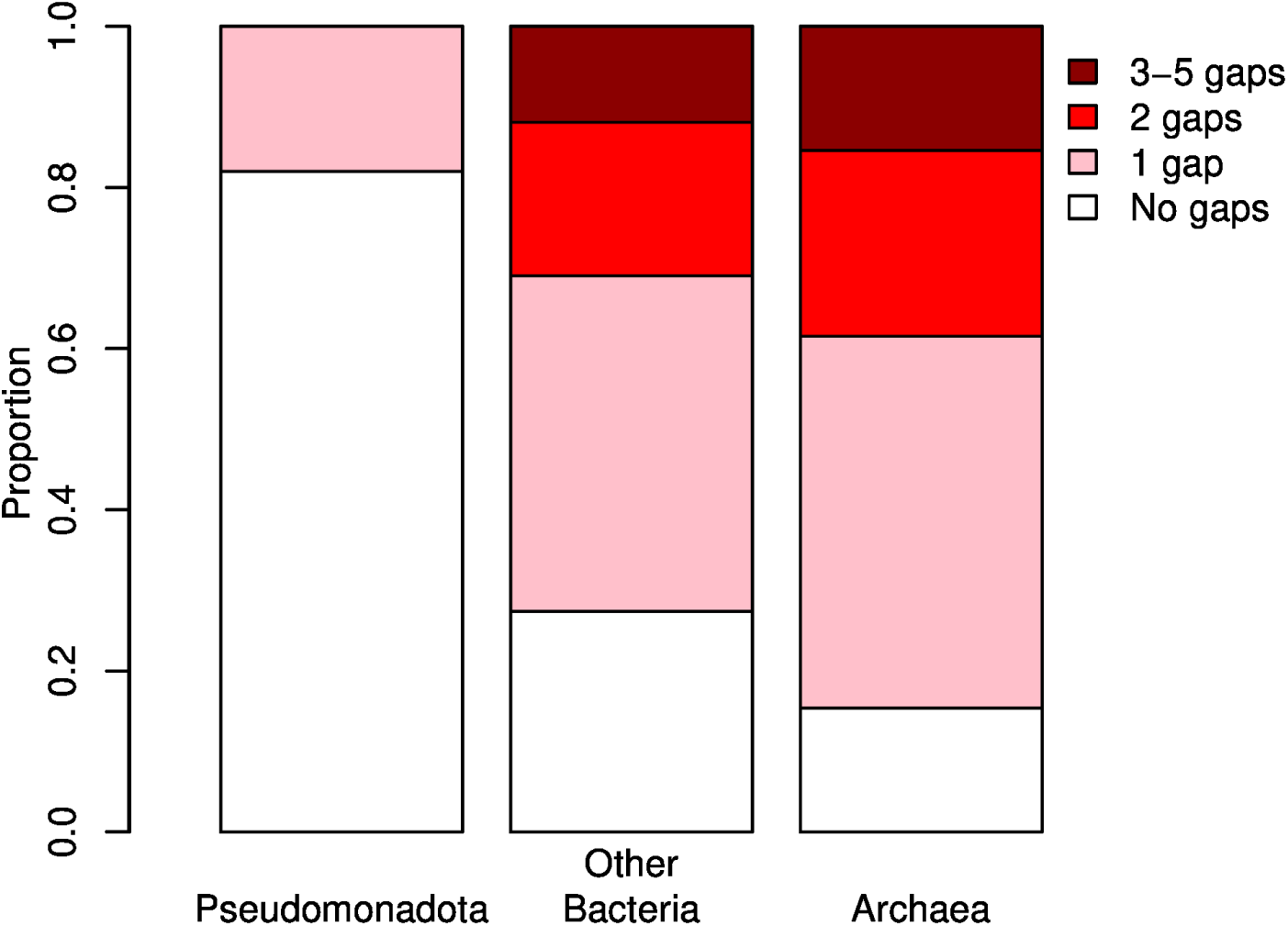
Genuine gaps in the amino acid biosynthesis pathways of prototrophic bacteria and archaea. Gaps that are due to missing gene calls or frameshifts (which may be spurious) were not included.

### The accuracy of GapMind for auxotrophic bacteria

We also tested GapMind on prokaryotes with known auxotrophies. We identified 26 prokaryotes (all bacteria) with experimentally-confirmed requirements for one or more amino acids (see Methods). These gave a total of 106 genome x pathway combinations for which the pathway was reported not to be functional. GapMind found that 102 of these (96%) had at least one gap or low-confidence step.

We manually examined the four cases where the GapMind results might incorrectly imply that the strain was prototrophic for that amino acid. Three of these cases involve *Streptococcus pneumoniae* D39. As previously reported, *S. pneumoniae* appears to contain all of the genes necessary for isoleucine and valine synthesis, and it is not clear why these amino acids are required for growth (Kazmierczak et al. 2009). Also, *S. pneumoniae* is auxotrophic for glycine even though it encodes a likely serine hydroxymethyltransferase (*glyA,* SPD_RS04890), which in most bacteria converts serine to glycine. As discussed above, it appears that *S. pneumoniae* does not synthesize or take up serine, and its GlyA functions in reverse, to convert glycine to serine (Härtel et al. 2012).

Finally, *Fusobacterium varium* ATCC 27725 requires methionine for growth (Resmer and White 2011). Although GapMind did not identify any low-confidence steps in methionine synthesis in *F. varium*, two steps were medium-confidence: homoserine dehydrogenase and cystathionine gamma-synthase (*metB*). The potential homoserine dehydrogenase (C4N18_RS12320) is only 39% identical to a characterized homoserine dehydrogenase, but the active-site residues are conserved and C4N18_RS12320 is encoded in an apparent operon with two other genes for homoserine biosynthesis (aspartate kinase and aspartate-semialdehyde dehydrogenase). So we expect that C4N18_RS12320 is truly a homoserine dehydrogenase. Furthermore, *F. varium* grows in the absence of threonine (Resmer and White 2011), which suggests that homoserine dehydrogenase (which is usually required for threonine synthesis) is present. For *metB*, GapMind identified two medium-confidence candidates, both of which have greater similarity to methionine gamma-lyases (73-77% identity) than to known MetB proteins (44-48%). Given the growth data for *F. varium*, these proteins probably lack MetB activity. Overall, if a bacterium was reported to require an amino acid for growth, GapMind identified at least one low-confidence step in that pathway 96% of the time.

### Conclusions

The revised GapMind for amino acid biosynthesis includes novel genes for the synthesis of arginine, glycine, lysine, methionine, serine, and branched-chain amino acids, as well as many diverged enzymes. Across prototrophic bacteria and archaea, 95% of pathways are complete, with no low-confidence steps, while for experimentally-identified auxotrophies, 96% of pathways had at least one low-confidence step. The majority of the remaining gaps in functioning pathways are due to just four steps: *hisN*, *hisI, serB,* and *thrB*.

## Materials and Methods

### Updating GapMind

Besides including the additional characterized enzymes and predictions described above, we also updated GapMind to use newer versions of the curated databases Swiss-Prot (UniProt Consortium 2023), MetaCyc (Caspi et al. 2020), BRENDA (Chang et al. 2021), and the Fitness Browser reannotations (Price et al. 2018b; Price and Arkin 2024b). These were updated in November 2023. (GapMind also uses CharProtDB, which, as far as we know, has not been updated (Madupu et al. 2012).)

GapMind for amino acid biosynthesis now includes 159 different biosynthetic steps. These are described by 2,261 characterized proteins from curated databases, 138 characterized proteins that we curated, 4,469 curated predictions from Swiss-Prot, and 30 proteins whose function we predicted. Candidates that were identified from their similarity to proteins with predicted functions -- either from the predictions described above, or from curated annotations in Swiss-Prot that lack experimental evidence -- are treated as one level of confidence lower than if they were based on similarity to a characterized protein. For instance, if a candidate is at least 40% identical to a protein with a predicted function, and the alignment covers at least 70% of the protein with a predicted function, then the candidate will be considered medium confidence. GapMind also uses hidden Markov models of protein families to describe some steps. These are based on 141 TIGRFams (Haft et al. 2013) and four PFams (Finn et al. 2014).

When run from the command-line, GapMind now provides the option to use DIAMOND (Buchfink et al. 2015) instead of UBLAST (Edgar 2010) for pairwise searches. DIAMOND is not as fast as UBLAST, but is more sensitive. Because GapMind ignores homologs that are less than 30% identity or that cover less than half of the characterized protein, this makes little difference to the final results of GapMind.

### Essential proteins

Essential proteins for *Echinicola vietnamensis* KMM 6221 (DSM 17526), *Pedobacter sp.* GW460-11-11-14-LB5, and *Pontibacter actiniarum* KMM 6156 (DSM 19842) were reported previously (Price et al. 2018b). Essential proteins were defined as proteins with unusually low coverage by transposon mutants (Price et al. 2018b). Mutants of these genes might grow very slowly, rather than being truly essential. A RB-TnSeq library for *Mucilaginibacter yixingensis* YX-36 was described elsewhere (Torres et al. 2025); from this library, 447 essential proteins were identified, using the same approach. Similarly, based on an RB-TnSeq library for *Lysobacter soli* OAE881 (see below), we identified 253 essential proteins.

### Sequence analysis

Characterized homologs were identified using PaperBLAST (Price and Arkin 2017). Functional residues were identified using SitesBLAST (Price and Arkin 2022). Conserved gene neighbors were identified using *fast.genomics* (Price and Arkin 2024a). RB-TnSeq data was examined using the Fitness Browser (Price and Arkin 2024b).

For finding homologs within GapMind, we used UBLAST (Edgar 2010) from 64-bit usearch 11.0.667 (available at https://github.com/rcedgar/usearch_old_binaries/) and HMMer 3.3.1 (http://hmmer.org/; (Eddy 2011)). Boltz-2x, which is the co-folding prediction tool Boltz-2 with a physical steering potential (Passaro et al. 2025), was invoked with a protein sequence and a SMILES string for the small molecule, using foldy (Roberts et al. 2024). The predicted structure of GlyXL from *Bifidobacterium breve* (bound to glyoxylate) was very similar to the experimental structure (PDB:2HA9) for GlyXL from *Streptococcus pneumoniae* (root mean square deviation of 1.21 Å, TM-score 0.97, 59% sequence identity), as assessed using RCSB (Bittrich et al. 2024).

### Prokaryotes that grow in defined media

We previously reported a list of 148 strains of prokaryotes that grow in minimal media and have complete genome sequences available ((Price et al. 2020); derived from (Dos Santos et al. 2012; Oberhardt et al. 2015)). One of those genomes (GCF_000195895.1, for *Methanosarcina barkeri* str. Fusaro) has been removed from Genbank, so we removed it from our list. We also reported a list of 35 bacteria that had RB-TnSeq data during growth in minimal media (Price et al. 2020). Recent studies reported RB-TnSeq data for four additional bacteria growing in minimal media: *Magnetospirillum magneticum* AMB-1 (McCausland et al. 2022), *Mucilaginibacter yixingensis* YX-36 (DSM 26809) (Torres et al. 2025), *Nitratidesulfovibrio vulgaris* Hildenborough (Trotter et al. 2023), and *Xanthomonas campestris pv. campestris* strain 8004 (Luneau et al. 2022). For this study, we conducted RB-TnSeq assays for *Brevundimonas vesicularis_C_* GW460-12-10-14-LB2, *Lysobacter soli* OAE881 (DSM 113522), and *Rhodanobacter denitrificans* FW104-10B01 (see below).

We identified additional prototrophic prokaryotes from the literature: *Thermoanaerobacter kivui* LKT-1 (Basen et al. 2018), *Thermus aquaticus* YT-1 (Brock and Freeze 1969), *Haloferax volcanii* DS2 (Trieselmann and Charlebois 1992), *Nitrosopumilus maritimus* SCM1 (Qin et al. 2017), *Streptomyces albulus_C_* NBRC14147 (Hasebe et al. 2024), and *Dissulfurispira thermophila* T55J (Umezawa et al. 2021). Also, a recent study identified several isolates from the human gut that are prototrophic (Soto-Martin et al. 2020); we included *Anaerobutyricum hallii* DSM 3353, *Roseburia faecis* M72, and *Clostridium tyrobutyricum* FAM22553. A caveat for these three strains is that cysteine was included in the growth medium as a reductant; however, cysteine synthesis appears to be complete in all three strains, so we expect that they are truly prototrophic. Similarly, cysteine was present in the growth medium for *T. kivui*, but since it is autotrophic (Basen et al. 2018), it is presumably able to make cysteine.

Finally, to increase the diversity of prokaryotes in our list, we used the IJSEM database (Barberán et al. 2017) to identify representatives of taxonomic orders that were not included in the list, but had been reported to utilize specific carbon sources. We then checked the species descriptions to see if they used a truly defined medium with no added amino acids. By this approach, we identified ten prototrophic type strains (*Algiphilus aromaticivorans* DG1253, *Aquimarina longa* SW024, *Carboxydothermus pertinax* Ug1, *Halococcus hamelinensis* 100A6, *Hippea alviniae* EP5-r, *Methanospirillum lacunae* Ki8-1, *Nocardioides dokdonensis* FR1436, *Phaeacidiphilus oryzae* TH49, *Thiohalospira halophila* DSM 15071 HL 3, and *Tistlia consotensis* USBA 355).

### Experimentally-determined amino acid auxotrophies

Most of the experimentally-determined auxotrophies were taken from previous reports (Ashniev et al. 2022; Ramoneda et al. 2023). We removed some species because the original reference did not identify which exact strain was auxotrophic, and we wanted to ensure that the genome sequence analyzed was from an auxotrophic strain. We removed some strains because the precise amino acids that were required for growth were not determined. We also identified several strains that were proposed to be auxotrophic for cysteine, but based on experiments with sulfate as the sole source of sulfur, and the pathway for sulfate assimilation is absent from the genome (i.e., (Hébert et al. 2004; Ferrario et al. 2015)). In these cases, reduced sulfur compounds such as sulfide may be the natural sulfur source, and if a suitable sulfur source was provided, cysteine synthesis may well occur. Finally, we removed *Klebsiella pneumoniae* KP11 because growth data was not reported and we removed *Yersinia ruckeri* YRB because there was significant residual growth after individual amino acids were removed from the media (Seif et al. 2020).

We added some additional auxotrophies from the literature: *Cysteiniphilum litorale* DSM 101832 is auxotrophic for cysteine (Liu et al. 2017); *Bacillus subtilis* 168 is auxotrophic for tryptophan (Zeigler et al. 2008); *Myxococcus xanthus* DK101 is auxotrophic for all three branched-chain amino acids (Bretscher and Kaiser 1978); *Kytococcus sedentarius* DSM 20547 is auxotrophic for methionine (Stackebrandt et al. 1995); *Legionella pneumophila* Philadelphia 3 is auxotrophic for arginine, methionine, serine, threonine, and valine (Ristroph et al. 1981); and *Streptococcus pneumoniae* D39 is auxotrophic for arginine, glycine, histidine, isoleucine, leucine, and valine (Kazmierczak et al. 2009).

In total, we had 26 auxotrophic bacteria, with 106 auxotrophies for amino acids. Four of these bacteria were auxotrophic for all three aromatic amino acids. In principle, these bacteria could be auxotrophic for chorismate (the common precursor for the aromatic amino acids) while being capable of converting chorismate to all three aromatic amino acids. This would complicate the comparison of the growth requirements to GapMind’s pathways, because GapMind describes chorismate synthesis separately from the biosynthetic pathways for phenylalanine, tyrosine, and tryptophan. In reality, all four of the bacteria have gaps in all three pathways downstream of chorismate. Two of the four have gaps in chorismate synthesis as well (*Lactobacillus delbrueckii subsp. lactis* CRL581 and *Lactobacillus paracasei* LC2W).

### RB-TnSeq libraries

The mutant libraries for *Brevundimonas vesicularis_C_* GW460-12-10-14-LB2 and *Bacteroides thetaiotaomicron* VPI-5482 were described previously (Liu et al. 2018).

To construct a mutant library in *Lysobacter soli* OAE881 (Coker et al. 2022), we used conjugation from *E. coli* WM3064 harboring the pHLL250 *mariner* transposon vector library (strain AMD290), which was previously built via Golden Gate assembly (Liu et al. 2018). Specifically, we grew 10 mL of wild-type OAE881 in LB overnight at 30°C. The next morning, we recovered a 2 mL freezer stock of strain AMD290 in 50 mL LB supplemented with 50 mg/mL carbenicillin (Cb) and 300 µM diaminopimelic acid (DAP) at 37°C. When the OD_600_ of the *E. coli* donor strain reached 1, we harvested 20 OD_600_ units of the culture and washed the cells three times with fresh LB supplemented with DAP. Then, 20 OD_600_ units of wild-type OAE881 cells were harvested, mixed with the washed donor cells, centrifuged, and resuspended in a final volume of 0.5 mL with LB supplemented with DAP. The resuspension was spotted onto 0.45-µm membrane filters (Millipore, United States) and incubated overnight on LB agar plates supplemented with DAP at 30°C. After 16 h, the conjugation mixture was scraped from the membrane, resuspended in 10 mL LB with 50 µg/mL kanamycin (Km) and plated at different dilutions on LB plates supplemented with 50 µg/mL Km.

Plates were incubated at 30°C for 48 h to let visible colonies develop. We then pooled the colonies and grew the library in liquid LB supplemented with 50 µg/mL Km for 2 population doublings. We then added glycerol to a final volume of 15%, made multiple 1 mL −80°C freezer stocks (∼10^8^ cells/mL) of the final library for subsequent experiments, and collected cell pellets to extract genomic DNA for TnSeq mapping. To map the genomic locations of the transposon insertions and link these insertions to their associated DNA barcodes, we used a variation of the previously described TnSeq protocol (Wetmore et al. 2015), where we use a splinkerette adaptor instead of a Y adapter and we used two rounds of PCR to selectively enrich for transposon junctions (Rubin et al. 2022). We mapped 118,245 barcodes to insertions in the genome.

To construct a mutant library in *Rhodanobacter sp.* FW510-R12, we followed the same conjugation procedure as described above for *Lysobacter soli* OAE881, except that mutants were selected on R2A with 5 µg/mL Km at 25°C. We mapped 478,711 barcodes to insertions in the genome of FW510-R12.

To construct a mutant library in *Rhodanobacter denitrificans* FW104-10B01, we first developed a custom mariner transposon delivery vector, pTKO49_NN1. pTKO49_NN1 was modified from our standard mariner vector (pHLL250_NN1, (Adler et al. 2021)), to avoid methylated motifs that were identified in the genome of FW104-10B01 from the PacBio sequencing data (Peng et al. 2022) and which may be targeted by restriction enzymes. These motifs were: GACCAG (removed 1 instance), CTCGAG (removed 1 instance), ATCNNNNNNTGGA (removed 2 instances), TCCANNNNNNGAT (removed 2 instances), GAANNNNNTTGG (removed 2 instances), TTCANNNNNCTC, and GAGNNNNNTGAA, where the last two motifs did not appear in pHLL250_NN1. Compared to pHLL250_NN1, pTKO49_NN1 has eight single nucleotide changes and was made in a two step process, using our previously described parts-based "Magic Pools" approach (Liu et al. 2018). First, we used Golden Gate assembly with five parts to make the non-barcoded vector pKTO49. The part1 vector (pTKO2) containing the majority of the mariner transposase gene was generated from gBlock gHL47 (Integrated DNA Technologies) and cloned into the pJW52 holding vector via Gibson assembly as previously described (Liu et al. 2018). The part2 vector pHLL216 (Liu et al. 2018) contains the kanamycin resistance gene promoter. The part3 vector (pTKO1) contains the kanamycin resistance gene and was generated from gBlock gHL48. The part4 vector pTKO22 contains the plasmid backbone, a region for incorporating DNA barcodes, a carbenicillin resistance gene, the colE1 origin of replication, and GFP. pTKO22 is two nucleotides different from the part4 in pHLL215 (Liu et al. 2018) and the two mutations were introduced by amplifying two parts of pHLL215, one with oAD474 and oAD475 (where the middle of oAD475 introduces the two mutations) and the second with oAD476 and oAD477; fusing these two PCR products via PCR with oAD477 and oAD478; and cloning this final PCR product into pJW52. The part5 vector pJW14 (Liu et al. 2018) contains the promoter for the transposase. pTKO49 was generated with BbsI-mediated Golden Gate assembly with the 5 part vectors as described (Liu et al. 2018) and was sequence verified. We then incorporated millions of DNA barcodes into pTKO49 to make pTKO49_NN1 using BsmBI-mediated Golden Gate assembly, as described (Liu et al. 2018). We moved the pKTO49_NN1 plasmid library into the *E. coli* conjugation donor WM3064 via electroporation to make the conjugation donor strain AMD1385. For primer sequences, see Supplementary Table S3; for gBlock sequences, see Supplementary Table S4; and for plasmids, see Supplementary Table S5.

The mutant library of FW104-10B01 was made by conjugation between recipient wild-type FW104-10B01 and strain AMD1385. An aliquot of the AMD1385 donor was grown overnight in 50 mL of LB supplemented with DAP and carbenicillin. To remove residual carbenicillin, we pelleted (via centrifugation) and washed the AMD1385 cells two times with LB supplemented with DAP only. These washed donor cells were mixed in a 1:1 ratio with wild-type FW104-10B01, and the combined cell mix was pelleted by centrifugation. We resuspended the combined pellet in 1 mL of LB with DAP, spread this cell slurry across 4 LB plates supplemented with DAP, and incubated these plates for 12 hours at 30°C. After conjugation, we harvested the cell mixture by scraping up the cells into 10 mL of R2A supplemented with 15% glycerol, and froze down 10 1-mL aliquots at -80°C. One of these conjugation aliquots was used to estimate the number of mutant colony forming units (CFU) in our conjugation mixture. Briefly, we thawed this tube, performed 10-fold serial dilution in R2A, plated these dilutions on R2A plates supplemented with 10 ug/mL kanamycin, incubated these plates for 4 days at 30°C, and counted the number of CFUs. Based on these numbers, we thawed additional aliquots of the conjugation mixture and plated across 84 R2A plates supplemented with 10 ug/mL kanamycin. After 4 days incubation at 30°C, we scraped up all of the mutant colonies into 10 mL of R2A, measured the OD_600_ of this mixture, and inoculated these cells into 100 mL fresh R2A with 10 ug/mL kanamycin at a starting OD_600_ of 0.25. The cells were grown for 12 hours at 30°C to allow a few population doublings, glycerol was added to a final concentration of 15%, and we made single-use aliquots of the final mutant library (rhodanobacter_10B01_ML12). We also extracted genomic DNA at this point for TnSeq mapping of the transposon insertions. We mapped 507,095 barcodes to insertions in the genome of FW104-10B01.

To construct a mutant library in *Rhodanobacter* sp. FW510-T8, we used conjugation with *E. coli* donor strain AMD1385 carrying pTKO49_NN1. Transconjugants were selected on R2A agar with 10 µg/mL kanamycin at 25°C and pooled to generate the FW510-T8 RB-TnSeq library. The final library was constructed by collecting colonies from approximately 150 selection plates, corresponding to an estimated >500,000 mutants. We mapped 719,225 barcodes to insertions in the genome of FW510-T8.

### RB-TnSeq assays

We performed fitness assays as described previously (Wetmore et al. 2015). Briefly, the mutant library was recovered from the freezer in the rich media that used to make the library, inoculated at OD_600_ = 0.02 into the condition of interest, grown until saturation, genomic DNA was extracted, and barcodes were amplified with PCR and sequenced using Illumina. Fitness values (log2 ratios) were calculated as described previously (Wetmore et al. 2015).

For *Brevundimonas vesicularis_C_* GW460-12-10-14-LB2, we performed experiments in a defined minimal medium at 20°C or 30°C with 20 mM glucose or 0.5% v/v Tween-20 as the carbon source. The basal medium (RCH2_defined_noCarbon) included 0.25 g/L Ammonium chloride, 0.1 g/L Potassium Chloride, 0.6 g/L Sodium phosphate monobasic monohydrate, 30 mM PIPES sesquisodium salt, Wolfe’s mineral mix (ATCC), and Wolfe’s vitamins (ATCC).

For *Lysobacter soli* OAE881, we performed fitness assays with 17 defined carbon sources (each at 20 mM) as well as casamino acids at 30°C and with 200 rpm shaking, again with RCH2_defined_noCarbon as the basal medium.

For *Rhodanobacter denitrificans* FW104-10B01, we performed a variety of stress experiments in R2A medium, which will be described elsewhere (H. Lesea et al, in preparation). For this study, we performed 30 fitness experiments in defined media. Two of these experiments were conducted in a minimal defined medium (RCH2_defined_noCarbon) with 20 mM glucose, at either 20°C or 30°C. For the remainder of the experiments, we used 13 different carbon sources, and we added 100 µM of each amino acid to the medium.

For *Rhodanobacter sp.* FW510-R12, we performed fitness assays in NLDM_defined_nocarbon medium with 20 mM glucose ± 1 mM L-serine at room temperature. (NLDM_defined_nocarbon includes 1 mM ammonium chloride, 3.3 mM potassium chloride, 0.812 mM magnesium sulfate, 0.68 mM calcium chloride, 4.05 mM disodium phosphate, 0.95 mM sodium phosphate monobasic, ATCC Wolfe’s mineral mix, and ATCC Wolfe’s vitamin mix.) Experiments were performed in glass test tubes (10 ml culture volume) and shaken at 200 rpm.

For *Rhodanobacter sp.* FW510-T8, we performed fitness assays in NLDM_defined_nocarbon medium with 20 mM glucose ± 1 mM L-methionine at room temperature. Experiments were performed in 96-well deep well blocks with a culture volume of 1 mL and shaken at 700 rpm.

For *Bacteroides thetaiotaomicron* VPI-5482, fitness assays were performed anaerobically in four different nitrogen sources using Varel Bryant medium (Liu et al., 2021). The basal medium was modified to minimize or eliminate nitrogen by replacing the L-methionine and L-cysteine with either (134 **μ**M L-methionine + 3mM L-cysteine) or (134 **μ**M L-methionine + 3mM dithiothreitol + 3mM sodium sulfide) or (0.1 ng/L cyanocobalamin + 3mM dithiothreitol + 3mM sodium sulfide).

### Cloning of putative biosynthetic genes

Candidate biosynthetic genes were cloned to replace RFP in pBbA2c-RFP (AddGene plasmid 35326; (Lee et al. 2011)), which contains a p15A origin, CmR for chloramphenicol resistance,the *tetR* repressor, and a *tet*-inducible promoter driving the expression of RFP. As previously described (Price et al. 2021), the pBbA2c-RFP vector was linearized by PCR and the target genes’ inserts were either PCR amplified from genomic DNA or ordered as gBlocks (Integrated DNA Technologies). Sequences for primers are in Supplementary Table S3; sources of genomic DNA for cloning are listed in Supplementary Table S6; and sequences for gBlocks are in Supplementary Table S4. After cloning using the Gibson Assembly Mastermix (New England Biolabs), plasmids with the correct sequence were identified by Sanger sequencing (Elim Biopharmaceuticals) and are listed in Supplementary Table S5.

### Complementation assays

Many of the complementation assays used strains of *E. coli* from the Keio collection in which a gene of interest (*thrB, glyA, serB, serC,* or *argE*) is replaced with a kanamycin resistance gene (Baba et al. 2006). To study the alternative *dapE* from *Echinicola* or *Pedobacter*, we used the AT978 strain of *E. coli* (Bukhari and Taylor 1971). AT978 was obtained from the Coli Genetic Stock Center and its genotype is LAM- e14- *dapE9 relA1 spoT1 thiE1*.

To construct the *dapF* knockout strain in *E. coli* strain MG1655, we used a modified CRISPR/Cas9 system described previously (Li et al. 2021). Briefly, the recipient strain containing the pEcCas plasmid encoding Cas9 and λ-Red recombinase is transformed by electroporation with the pEcgRNA plasmid carrying a dapF-specific sgRNA and a specific linear DNA template. The specific N20 spacer was predicted using the CCTop website (https://cctop.cos.uni-heidelberg.de/) (Stemmer et al. 2015; Labuhn et al. 2018).The knockout DNA template was created by combining the upstream and downstream region of *dapF* using overlapping PCR. The transformants were selected on kanamycin and spectinomycin (both at 50 μg/mL). After verification by PCR and sequencing, the positive clones were cured of both plasmids.

For all the strains used in the complementation assays, plasmids were introduced in the appropriate strains by electroporation and transformants were selected on LB supplemented with 20 μg/mL chloramphenicol.

Growth assays were run in 96-well plate format in either Tecan Sunrise or Agilent Epoch2 plate readers. The temperature ranged from 27°C to 30°C and the shaking speed from ∼300 to 550 rpm depending on the instrument. All strains for complementation assays (Table 5) were grown in minimal M9 medium with 20 mM glucose and 20 µg/mL chloramphenicol to maintain the plasmid. We sometimes added 4 nM of anhydrotetracycline (aTc) as an inducer. We found that aTc at this concentration moderately inhibited the growth of *E. coli* carrying the control plasmid (which expresses RFP), and in general, strains complemented with the *E. coli* gene grew better without aTc, but aTc was often needed for good growth of strains with heterologous complementation.

**Table 5:** Strains for complementation assays.

| Strain | Background | Plasmid | Description |
| --- | --- | --- | --- |
| AMD806 | BW25113 | pBbA2c-RFP | wild-type with RFP expressed from pBbA2c |
| AMD3424 | BW25113 *thrB::kan* | pBbA2c-RFP | Δ*thrB* with pBbA2c-RFP |
| AMD3425 | BW25113 *thrB::kan* | pALF4 | Δ*thrB* with BT2402 cloned into pBbA2c |
| AMD3426 | BW25113 *thrB::kan* | pALF5 | Δ*thrB* with DET_R08S350 cloned into pBbA2c |
| AMD3427 | BW25113 *thrB::kan* | pALF17 | Δ*thrB* with *E. coli thrB* cloned into pBbA2c |
| AMD3428 | BW25113 *glyA::kan* | pBbA2c-RFP | Δ*glyA* with pBbA2c-RFP |
| AMD3429 | BW25113 *glyA::kan* | pALF1 | Δ*glyA* with BBR_RS12915-BBR_RS12920 cloned into pBbA2c |
| AMD3431 | BW25113 *glyA::kan* | pALF15 | Δ*glyA* with *E. coli glyA* cloned into pBbA2c |
| AMD3432 | BW25113 *serB::kan* | pBbA2c-RFP | Δ*serB* with pBbA2c-RFP |
| AMD3433 | BW25113 *serB::kan* | pALF3 | Δ*serB* with CBO1128 cloned into pBbA2c |
| AMD3434 | BW25113 *serB::kan* | pALF16 | Δ*serB* with *E. coli serB* cloned into pBbA2c |
| AMD3451 | BW25113 *serC::kan* | pBbA2c-RFP | Δ*serC* with pBbA2c-RFP |
| AMD3452 | BW25113 *serC::kan* | pALF7 | Δ*serC* with CBO1126 cloned into pBbA2c |
| AMD3453 | BW25113 *serC::kan* | pALF8 | Δ*serC* with A0A843E9R6 cloned into pBbA2c |
| AMD3454 | BW25113 *serC::kan* | pALF9 | Δ*serC* with A0A1T4W7T3 cloned into pBbA2c |
| AMD3455 | BW25113 *serC::kan* | pALF19 | Δ*serC* with *E. coli serC* cloned into pBbA2c |
| AMD3456 | BW25113 *argE::kan* | pBbA2c-RFP | Δ*argE* with pBbA2c-RFP |
| AMD3457 | BW25113 *argE::kan* | pALF6 | Δ*argE* with W3Y6L2 cloned into pBbA2c |
| AMD3458 | BW25113 *argE::kan* | pALF18 | Δ*argE* with *E. coli argE* cloned into pBbA2c |
| AMD3644 | MG1655 | pBbA2c-RFP | wild-type with RFP expressed from pBbA2c |
| AMD3442 | MG1655 Δ*dapF* | pBbA2c-RFP | Δ*dapF* with RFP expressed from pBbA2c |
| AMD3443 | MG1655 Δ*dapF* | pALF11 | Δ*dapF* with HMPREF2922_03580 cloned into pBbA2c |
| AMD3444 | MG1655 Δ*dapF* | pALF21 | Δ*dapF* with *E. coli dapF* cloned into pJB00059 |
| AMD3914 | AT978 | pBbA2c-RFP | AT978 (*dapE9*) with RFP expressed from pBbA2c |
| AMD3917 | AT978 | pALF23 | AT978 with *E. coli dapE* cloned into pBbA2c |
| AMD3916 | AT978 | pALF14 | AT978 with Echvi_1427 cloned into pBbA2c |
| AMD3915 | AT978 | pALF13 | AT978 with CA265_RS23900 cloned into pBbA2c |

For W3Y6L2 from *Veillonella denticariosi* cloned into an *argE::kan* strain of *E. coli*, our standard aerobic assay did not yield any growth. We were unsure about the metal cofactor of W3Y6L2, and M9 medium does not contain any added metals except magnesium. (We used M9 salts from BD Difco (provided at 5x), which contains 33.9 g/L anhydrous disodium phosphate; sodium phosphate often contains impurities of iron and other transition metals, which are required for the growth of wild-type *E. coli* (Soma et al. 2023).) We tried adding Wolfe’s minerals, but the strain complemented with W3Y6L2 still did not grow aerobically. For anaerobic assays, we used a plate reader housed in a Coy anaerobic chamber (90% N_2_, 5% CO_2_ and 2.5% H_2_ atmosphere) with no shaking.

### Analysis of diaminopimelate secretion

Various strains of *E. coli* were grown in M9 glucose medium containing 20 µg/mL chloramphenicol, with or without 4 nM of aTc, for 16 hours at 30°C. Cultures were centrifuged at 7,000 rpm to remove cells and supernatants were filtered (0.22um PVDF), then lyophilized until dry. Dried materials were stored at -80°C until just prior to analysis, and then resuspended in 150µL of 100% methanol containing internal standards (Dataset S1). Samples were vortexed 2x10 seconds, sonicated in an ice water bath for 15 minutes, centrifuged (10,000 rcf, 5 min, 10°C), filtered (0.22um PVDF) and then filtrates were transferred to amber glass vials for LC-MS/MS analysis. Metabolites were separated using hydrophilic interaction liquid chromatography (HILIC) for polar metabolomics. Analyses were performed using an InfinityLab Poroshell 120 HILIC-Z column (Agilent) on an Agilent 1290 HPLC connected to a Q Exactive hybrid quadrupole-Orbitrap mass spectrometer (Thermo Fisher Scientific) using ElectroSpray Ionization (ESI). LC-MS/MS methods were previously described (B. Louie et al. 2024); run specific parameters are provided in Dataset S1. Peak heights and MS2 were extracted using mzmine 2.53 (Katajamaa et al. 2006); intensities and annotation evidence are provided in Dataset S1. Raw data files are available at GNPS2 (negative mode: https://gnps2.org/status?task=64d1b4b524674e42870ee88a4d6d8d6e; positive mode: https://gnps2.org/status?task=1398218c5d1e43909830a4b7c92b9c61)

## Availability of data and code

The code for GapMind, including the rule definitions, are available as part of the PaperBLAST code base (https://github.com/morgannprice/PaperBLAST). The code, the rules and the compiled rule definitions are archived at figshare (http://dx.doi.org/10.6084/m9.figshare.32680566). The figshare also includes a table of the 208 prototrophic bacteria and archaea, a table of the 26 bacteria with known auxotrophies, and the GapMind results for all of these genomes. The RB-TnSeq data is available from the Fitness Browser (http://fit.genomics.lbl.gov) and is archived at figshare (https://doi.org/10.6084/m9.figshare.32865896.v1).

## Supplementary Material

Supplementary tables S1-S6 http://morgannprice.org/GapMind/GapMind2026_Supplementary.xlsx

Dataset S1 http://morgannprice.org/GapMind/GapMind2026_DatasetS1.xlsx

## Appendix 1 Roles of *serB* and *serBK* in *Bacteroides thetaiotaomicron*

As part of a study of nitrogen source utilization by *Bacteroides thetaiotaomicron* VPI-5482, we collected fitness data with three amino acids as the sole sources of nitrogen: L-serine, L-asparagine, and L-glutamine. We found that *serBK* from *B. thetaiotaomicron* (BT1151; DUF1015 family) was important for fitness when L-serine was the nitrogen source (Figure S1). In previously-published experiments (Liu et al. 2021), *serBK* was also somewhat important for the utilization of D-serine, which is not explained.

**Figure S1:**
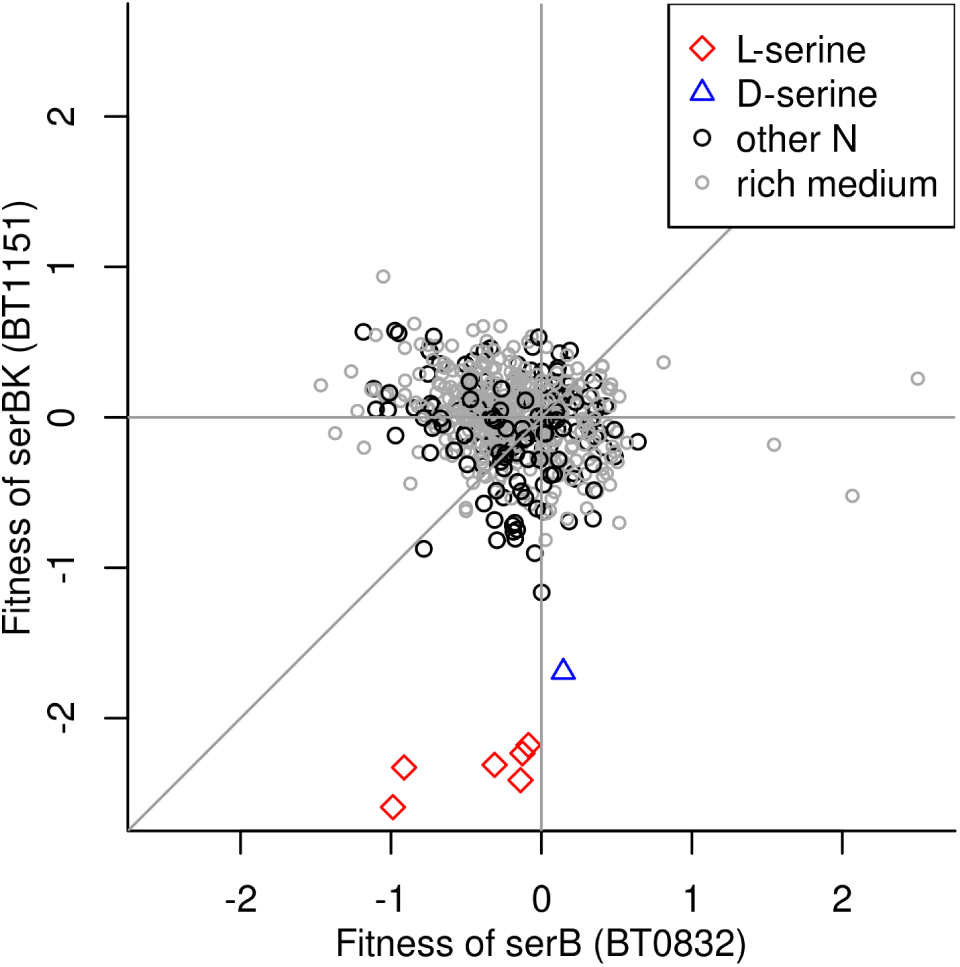
Fitness of serB and serBK in *Bacteroides thetaiotaomicron*. We highlight experiments with L-serine or D-serine as the sole source of nitrogen. Experiments with a defined medium with another nitrogen source (usually ammonium) are in black and experiments with a rich medium are in grey. Most of the data is from (Liu et al. 2021) and mouse experiments are not shown.

In most defined media conditions, neither *serB* (BT0832) nor *serBK* was important for fitness (Figure S1). It appears that either gene suffices for converting O-phospho-L-serine to L-serine.

## Appendix 2 Rationale for a putative alternative N-succinyl-L-glutamate synthase

*Steroidobacter denitrificans* DSM 18526 was reported to grow in defined minimal medium ((Fahrbach et al. 2008); DSMZ medium 1116), but its pathway for synthesizing arginine appears to be incomplete. In particular, from its genome, we could not identify any proteins with similarity to characterized N-acyl-L-glutamate synthases (ArgA). However, its genome does encode a cluster of genes for arginine synthesis, which includes a hypothetical protein (ACG33_RS14135; UniProt A0A127FCT3). Homologs of this hypothetical protein are virtually always found in arginine synthesis operons (Figure S2), which suggests a role in arginine synthesis. *S. denitrificans* is also missing the expected N-acylcitrulline hydrolase (ArgE), but some homologs of ACG33_RS14135 are found near plausible candidates for *argE* (purple genes in Figure S2). So we propose that ACG33_RS14135 is the missing ArgA.

**Figure S2:**
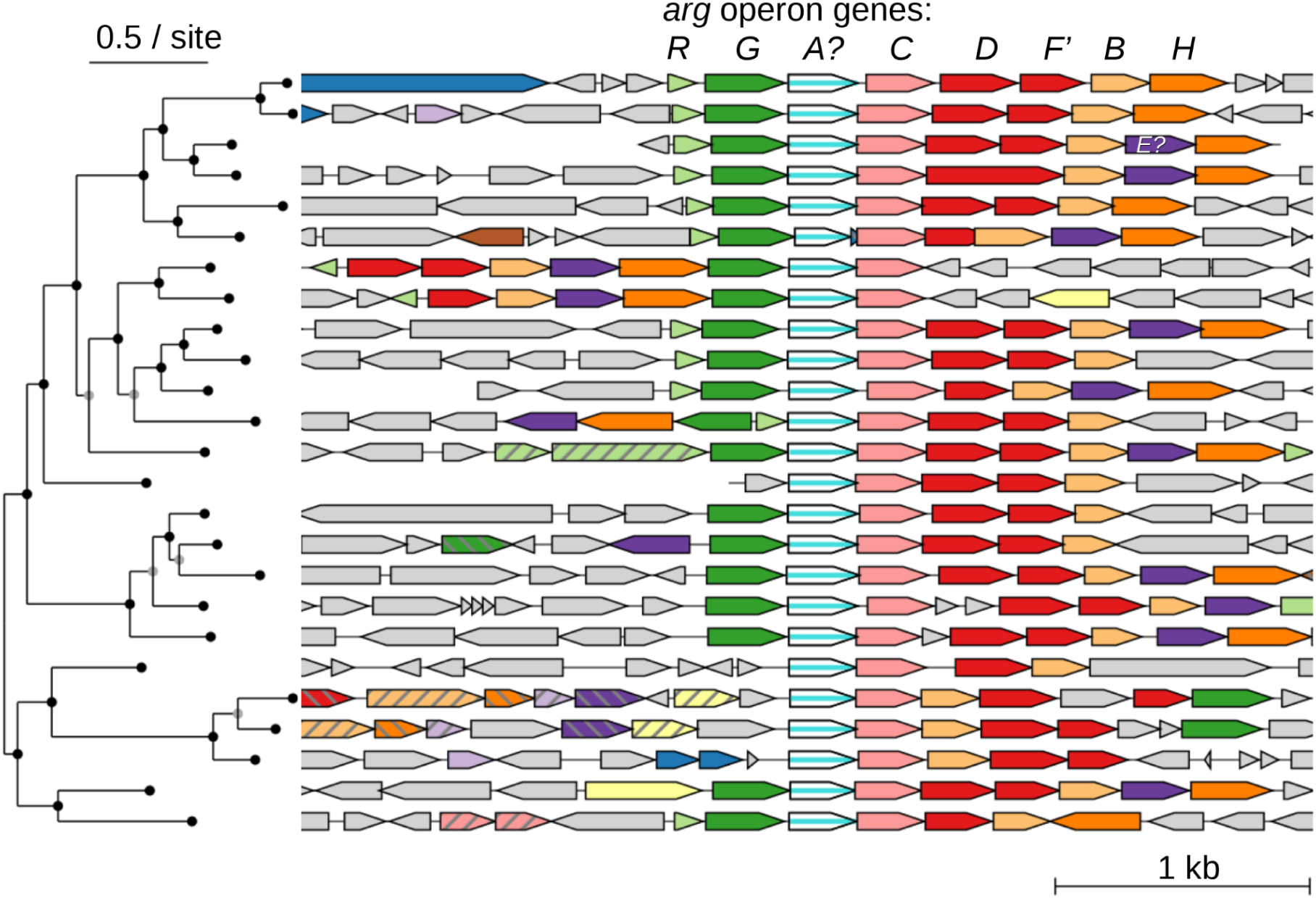
Gene neighbors for a putative alternate N-succinylglutamate synthase. Each row shows a homolog of the putative alternate *argA* (central column) from a different genus, along with its nearby genes. Other protein-coding genes are color-coded by sequence similarity. This figure was created using *fast.genomics* (Price and Arkin 2024a) and the central column includes 25 random proteins whose similarity to ACG33_RS14135 is at least 30% of the maximum bit score. The phylogenetic tree (at left) is based on the protein sequences of its homologs. Besides the proposed alternate *argA* (labeled *A?*), the genes labeled are: N-acylglutamate kinase *argB*; N-acyl-γ-glutamyl phosphate reductase *argC*; N-acylornithine aminotransferase *argD*; N-succinylornithine transcarbamoylase *argF*’; N-acylcitrulline hydrolase *argE* (in some of the clusters, in purple); argininosuccinate synthetase *argG*; argininosuccinate lyase *argH*; and transcriptional regulator *argR*.

If ACG33_RS14135 is replacing the usual ArgA, then homologs of both proteins should not be found in the same genome. We compared the distribution of homologs of ACG33_RS14135 to that of homologs of ArgA from *Pseudomonas aeruginosa* (AT700_RS27105), using *fast.genomics* (Price and Arkin 2024a). Potential functional orthologs (above 25% of the maximum bit score) were never found together in the same genome.

Using Foldseek (van Kempen et al. 2024), we found weak similarity between the predicted structure of ACG33_RS14135 (UniProt A0A127FCT3) and the experimentally-determined structure of a D-glutamate acetyltransferase ((Yu et al. 2023); PDB:7XRJ). Although this homology is remote (root mean square deviation = 5.6 Å, e-value = 1.4 · 10^-7^), it supports our proposal.

Although most bacteria synthesize arginine via N-acetylated intermediates, some bacteria use N-succinylated intermediates (Shi et al. 2006; Luque 2010). A single residue can determine whether the N-acylornithine transcarbamoylase (ArgF’) acts on N-succinylornithine or N-acetylornithine: mutating position 92 of an N-acetylornithine carbamoyltransferase from glutamate to alanine, proline, serine, or valine switches its substrate specificity to N-succinylornithine (Shi et al. 2007). Among 70 ArgF’ proteins encoded near ACG33_RS14135 or its homologs (from the *fast.genomics* database), 90% have A, P, S, or V at this position. If we also allow the chemically similar residues isoleucine or threonine, the proportion rises to 97%. (The two remaining homologs have a glutamine at this position.) So it appears that these organisms synthesize arginine via N-succinylated intermediates, and hence that these putative ArgA enzymes form N-succinylglutamate, not N-acetylglutamate. ACG33_RS14135 is included in the revised GapMind as a predicted ArgA.

## Appendix 3 Homoserine phosphotransferase HomK

As discussed in the main text, BT2402 is an alternative to homoserine kinase (BT2402) that is required for threonine synthesis in *Bacteroides thetaiotaomicron* (Price et al. 2020). BT2402 rescued the growth of a *thrB::kan* strain of *E. coli* in minimal medium, which confirms that BT2402 and related proteins (also known as TIGR02535 (Haft et al. 2013) or ThrB2 (Ashniev et al. 2022)) form O-phosphohomoserine. We named these proteins HomK. Close homologs of BT2402 (over 80% identical) from two species of a related genus, *Phocaeicola,* also seem to be involved in threonine synthesis. Specifically, both homologs have similar fitness patterns as threonine synthase, with cofitness = 0.88 in *P. dorei* CL03T12C01 (S. Tripathi and A.M. Deutschbauer, in preparation) and cofitness *=* 0.79 in *P. vulgatus* CL09T03C04 (data of (Huang et al. 2024)).

By analyzing the predicted structure of BT2402, we identified a likely specificity-determining residue that distinguishes HomK from homologs that are cofactor-independent phosphoglycerate mutases. Specifically, we identified the catalytic serine in BT2402 (S56) by aligning it to the phosphoglycerate mutase SSO0417 (Potters et al. 2003). This pxhosphorylated serine is thought to be an intermediate in the transfer of phosphate between substrates. When we folded BT2402 with phosphorylated S56 together with L-homoserine and two Mg^2+^ ions using Boltz-2 (Passaro et al. 2025), the phosphate group was placed near the hydroxyl group of homoserine, and the carboxylate side chain of a glutamate residue (E299) was just 2.8 Å from the amino group of homoserine (Figure S3A). As shown in Figure S3B, this glutamate is conserved in putative HomK proteins, while phosphoglycerate mutases almost always have a lysine at this position (consistent with a preference for negatively charged substrates).

**Figure S3:**
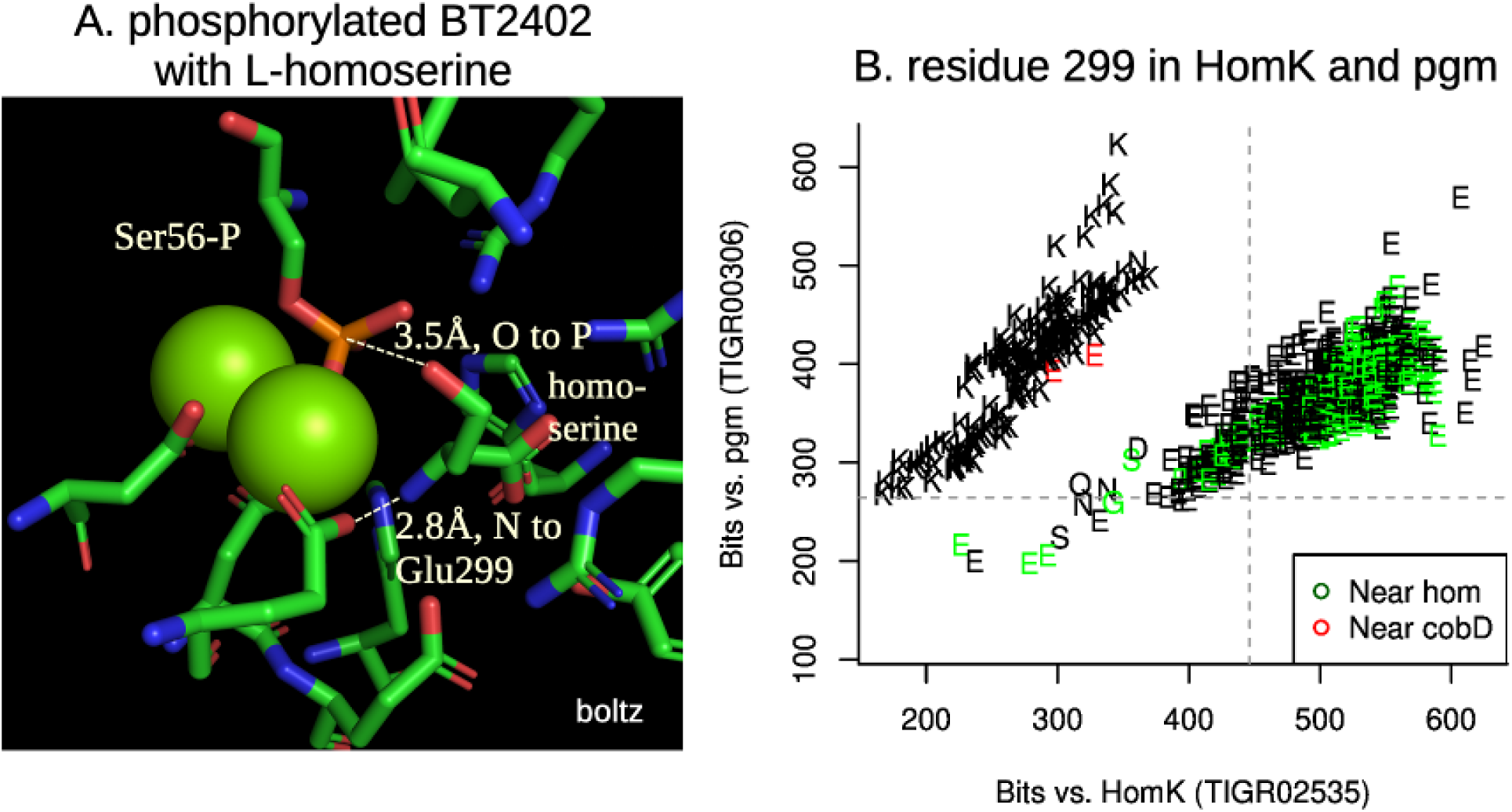
A putative specificity-determining residue differentiates HomK from cofactor-independent phosphoglycerate mutases. (A) Predicted model of BT2402 (HomK from *B. thetaiotaomicron*), with its catalytic serine 56 phosphorylated, bound to two Mg^2+^ ions (green spheres) and L-homoserine. (B) The putative specificity-determining residue (E299 in BT2402) across 1,012 homologs of BT2402 (30% amino acid identity or above and at least 300 a.a. aligning) from representatives of different genera (from *fast.genomics*). The *x* and *y* axes show how similar each protein is to the hidden Markov models for HomK (TIGR02535) or cofactor-independent phosphoglycerate mutases (TIGR00306) from TIGRFams (Haft et al. 2013), and the dotted lines show the “trusted” bit score cutoffs from TIGRFams. The plotting symbols show the specificity-determining residue. Colors highlight homologs that are encoded near and on the same strand as either homoserine dehydrogenase (*hom*), which forms the substrate of HomK, or *cobD*.

We did find two homologs of HomK that are more similar to the TIGRFam model for cofactor-independent phosphoglycerate mutases than to the model for HomK, yet have a glutamate aligning to E299. These are FWH37_03765 and G4O51_00460, both from metagenome-assembled genomes of Bathycorpusculaceae. These genes are encoded adjacent to L-threonine O-3-phosphate decarboxylase (*cobD)*, which is part of cobalamin synthesis. We speculate that FWH37_03765 and G4O51_00460 form L-threonine O-3-phosphate.

Among the homologs of BT2402 that are encoded near homoserine dehydrogenase and have conserved E299, we annotated several diverged homologs (41% identity or below) in GapMind. We verified that these homologs have conserved the other functional residues, namely the catalytic serine and the metal-coordinating residues. Cofactor-independent phosphoglycerate mutases rely on divalent metal cations for activity (van der Oost et al. 2002; Potters et al. 2003).

The experimental structure of the enzyme from *Pyrococcus horikoshii* (PDB:2zkt) shows two zinc ions, while the enzymes from *Sulfolobus* or *Pyrococcus* prefer magnesium, cobalt, or manganese. The coordinating residues in BT2402 would be D9, S56 (the catalytic serine), D346, and H347 for one metal ion and D274, H278, and H325 for the other. One of the diverged HomK proteins that we identified is DET_RS08350, which was confirmed by a complementation assay (see Appendix 5). The other diverged HomK proteins that we identified (UniProt:A0A2V1IX31, DCC48_11680, and UniProt:T2GDP6) are 33-41% identical to BT2402 or DET_RS08350 and are included as predictions in the revised GapMind.

What is the phosphate donor for HomK? Based on the homology to phosphoglycerate mutases, the most obvious donor would be 2- or 3-phosphoglycerate. In this case, the products of HomK would be O-phosphohomoserine and glycerate, and so we would expect that a glycerate kinase would be needed for the glycerate to reenter metabolism. We used AnnoTree (Mendler et al. 2019) to test the hypothesis that HomK (TIGR02535) co-occurs with glycerate kinases (KEGG ortholog groups K11529, K00865, or K15918). We found that only 53% of genomes that encode HomK encode one of the known forms of glycerate kinase. For comparison, 67% of genomes that encode that standard form of homoserine kinase (K00872; ThrB in *E. coli*) encode glycerate kinase. This suggests that phosphoglycerate is not the phosphate donor for HomK. Based on the putative specificity-determining residue E299, which should select for positively-charged substrates, we speculate that the donor is an amino acid such as phosphoserine. In fact, a phosphoserine:homoserine phosphotransferase (ThrH, not homologous to HomK) is able to substitute for ThrB in *Pseudomonas aeruginosa* (Singh et al. 2004).

## Appendix 4 Rationales for other previously-proposed alternate enzymes

The revised GapMind includes the thiocarboxylate pathway for converting O-phospho-homoserine to homocysteine, which was recently described in *Streptomyces albulus_C_* (Hasebe et al. 2024). The new steps are: sulfur carrier protein MetO; [MetO sulfur-carrier protein]-L-cysteine hydrolase Mec, which is required for the maturation of the N terminus of MetO; [MetO sulfur carrier protein]--sulfur ligase MoeZ; and MetO-thiocarboxylate-dependent L-homocysteine synthase MetM. Note that MetM is homologous to threonine synthases and is often misannotated as such.

*Methanocaldococcus jannaschii* synthesizes proline from serine via ornithine cyclodeaminase, but the protein responsible had not been identified (Graupner and White 2001). A recent study identified a novel family of ornithine cyclodeaminases in *Anabaena* and in *Methanococcus maripaludis* (Burnat et al. 2019). This family is present in *M. jannaschii* as well. Furthermore, the enzyme from *M. maripaludis* (UniProt Q6LXX7) is important for fitness unless proline is provided (data of (Day et al. 2024)). The revised GapMind includes these ornithine cyclodeaminases.

Ashniev and colleagues associated a family of putative transaminases with serine synthesis, via genome context, and named this family SerC2 (Ashniev et al. 2022). A mutant of SAUSA300_1669, which is a member of this family, is a serine auxotroph (Verstraete et al. 2018). This confirms that SerC2 is a family of phosphoserine transaminases.

As discussed in the main text, Ashniev and colleagues associated a family of putative transaminases with serine synthesis, which they named SerC3 (Ashniev et al. 2022), and we confirmed this using a complementation assay.

As discussed in the main text, Ashniev and colleagues associated a family of putative epimerases, which they named DapF2, with lysine synthesis (Ashniev et al. 2022). To test this, we used liquid chromatography-tandem mass spectrometry (LC-MS/MS) to measure the amount of diaminopimelate in the medium after the growth of four strains of *E. coli* in M9 glucose medium, with or without added inducer (aTc). These strains were: wild-type (*E. coli* MG1655) with a control plasmid (expressing RFP); a clean deletion of *dapF* in the MG1655 background (see Methods), also with the control plasmid; Δ*dapF* with *E. coli dapF* expressed on a plasmid; and ΔdapF with *dapF2* from *S. epidermis* (UniProt:A0A9Q5JK70, HMPREF2922_03580) expressed on a plasmid. As previously reported, we found that cells that lack *dapF* secrete diaminopimelate (DAP) into the medium (Mengin-Lecreulx et al. 1988). (Mengin-Lecreulx and colleagues reported that L,L-DAP, which is the substrate of DapF, was secreted, while our method did not distinguish between L,L-DAP or *meso*-DAP.) Furthermore, complementation with *dapF2* yielded similar results as wild-type cells (Figure S4). In the absence of inducer, complementation with *dapF* from *E. coli* yielded similar results as wild-type, but in the presence of inducer, expression of *dapF* from a plasmid yielded intermediate results; we speculate that this results from overly high expression of DapF. Overall, this experiment confirmed that *dapF2* can complement a *dapF* mutant, and hence that DapF2 is a diaminopimelate epimerase.

**Figure S4:**
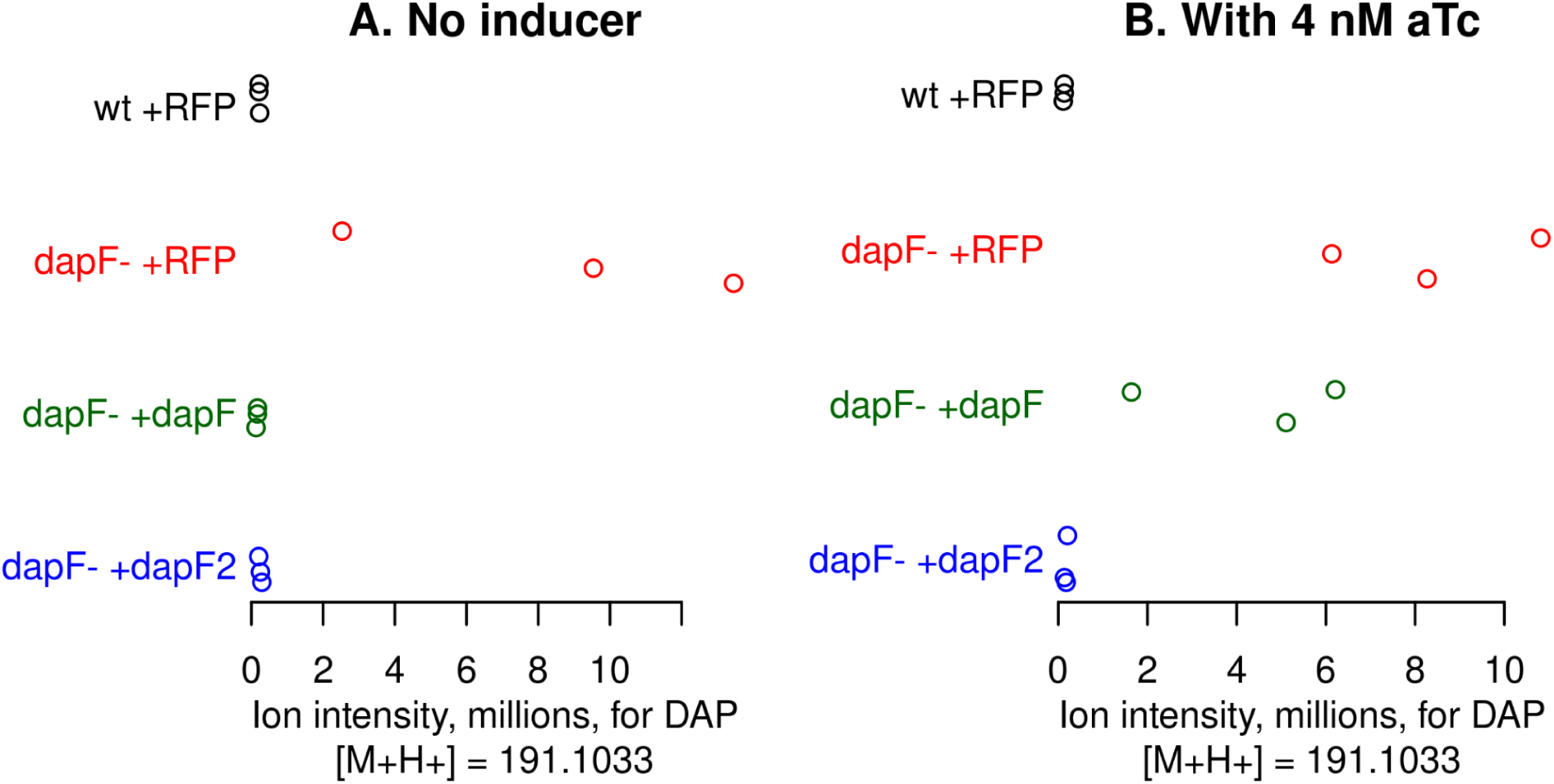
DapF2 from *Staphylococcus epidermis* prevents the secretion of diaminopimelate by *dapF* mutants of *E. coli*. We analyzed the supernatant of various strains of *E. coli* after growth in (A) M9 glucose medium or (B) M9 glucose medium with added inducer.

DapF2 is in a conserved operon with putative amidohydrolases, such as AA076_RS07060 (UniProt Q2FH40). These amidohydrolases have been proposed to be the missing N-acetyl-diaminopimelate deacetylase DapL (Jiang et al. 2015; Ashniev et al. 2022). AA076_RS07060 is 35% identical to N-acetyl-L-cysteine deacetylase SndA, which acts on a similar substrate, and has conserved catalytic residues.

We previously proposed several novel families of methionine synthases, including split MetE (cobalamin-independent synthase) and corrinoid-protein-dependent methionine synthase MesC (Price et al. 2021).

## Appendix 5 Diverged enzymes with experimental evidence

Eight of these enzymes were previously published. HisN in *Bifidobacterium breve* was recently identified by using RB-TnSeq (Shiver et al. 2025). TK0276 from *Thermococcus kodakarensis* is LysZ (LysW-glutamate kinase and LysW-2- aminoadipate 6-kinase) (Yoshida et al. 2016). Another six enzymes were confirmed using large-scale complementation assays (Biggs et al. 2024).

Seven enzymes were identified using fitness assays. First, two diverged enzymes in *Brevundimonas vesicularis_C_* GW460-12-10-14-LB2 were confirmed by conducting RB-TnSeq assays in minimal medium (see Methods): A4249_RS00005 is a diverged homoserine dehydrogenase and A4249_RS06835 is a diverged prephenate dehydratase.

Second, for the missing phosphoserine phosphatase (SerB) from *Lysobacter soli* OAE881 or *Xanthomonas campestris* 8004, we identified proteins that are similar to the biochemically characterized SerB from *Synechocystis* sp. PCC 6803 (Klemke et al. 2015). The genes from *L. soli* and *X. campestris* are in conserved operons with *serB* and both appear, based on RB-TnSeq data, to be essential ((Luneau et al. 2022); see Methods). These genes must encode the missing *serB*.

Third, acetolactate synthase / acetohydroxybutanoate synthase (AHAS) is a bifunctional enzyme that is involved in the biosynthesis of both valine and isoleucine. AHAS has a catalytic subunit and a regulatory subunit; the regulatory subunit is not strictly required for activity, but the catalytic subunit has far less activity on its own (Vyazmensky et al. 2009). The regulatory subunit (IlvH) usually has two domains, an ACT domain and an ACT-like domain, but some strains of *E. coli* encode an isozyme of AHAS whose regulatory subunit (IlvM) has a single ACT domain (Vyazmensky et al. 2009).

We noticed that some genera of Xanthomonadales that grow in minimal media lack the two-domain form of the regulatory subunit. Instead, we found proteins that were encoded near the catalytic domain but have only a single ACT domain, and are distantly related to *E.coli* IlvM (27-30% identity). To test the roles of these proteins, we used RB-TnSeq data for *Rhodanobacter denitrificans* FW104-10B01, *Lysobacter soli* OAE881, and *Xanthomonas campestris* pathovar *campestris* str. 8004 ((Luneau et al. 2022); see Methods). We found that in all three cases, mutants of the short *ilvM*-like gene (LRK54_RS10305, ABIE51_RS17555, or Xcc-8004.1058.1) had similar phenotypes as mutants in the adjacently-encoded catalytic subunit (Figure S5). This confirms that these are regulatory subunits.

**Figure S5:**
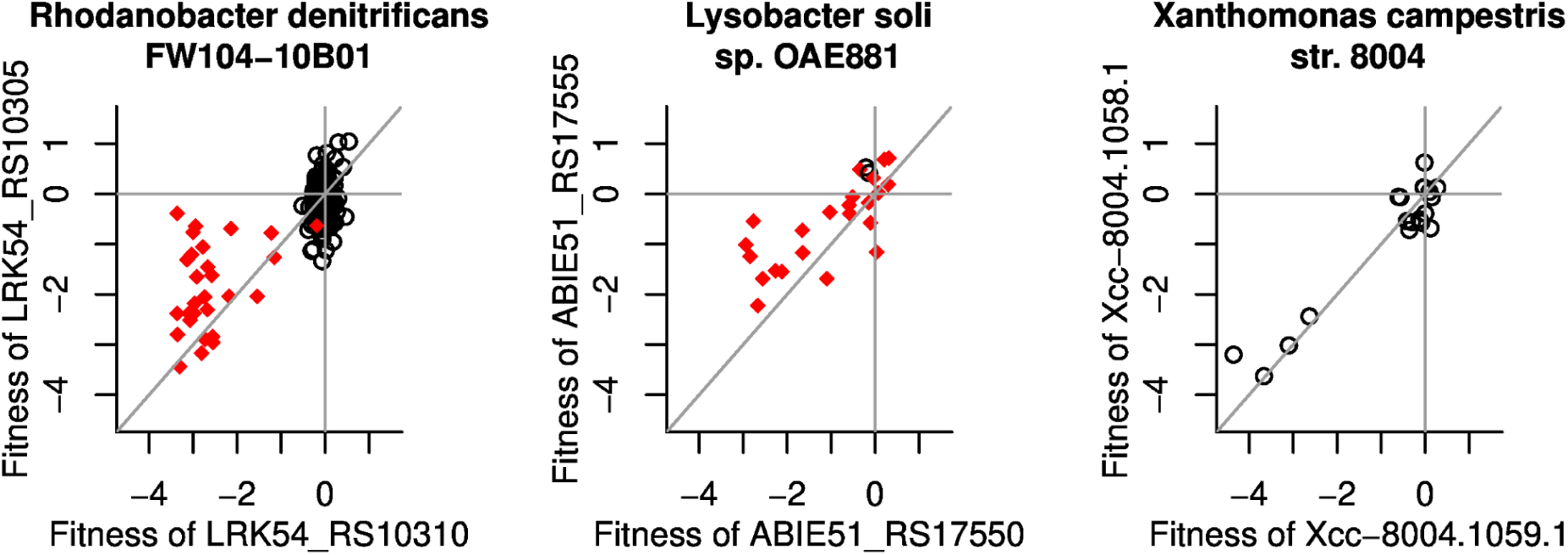
The putative short regulatory subunit of acetolactate synthase is cofit with the catalytic subunit in three different genera of Xanthomonadales. Fitness values (log2 ratios) are shown for the catalytic subunit (*x* axis) and the putative regulatory subunit (*y* axis). For *L. soli*, experiments in a defined minimal medium are highlighted in red. For *R. denitrificans*, experiments in a defined medium are highlighted; for many of these experiments, 100 μM of each amino acid was added. In each panel, the lines show *x* = 0, *y* = 0, and *x* = *y*.

Three additional enzymes were confirmed using complementation assays. First, we identified a divergent homolog of the homoserine phosphotransferase BT4202 (HomK) in *Dehalococcoides mccartyi*. DET_RS08350 is 39% identical to BT2402, but is classified by TIGRFam as phosphoglycerate mutase (TIGR00306) instead of as HomK (TIGR02535). Since *D. mccartyi* lacks *thrB* and encodes another phosphoglycerate mutase that is more distantly related to BT2402, we predicted that DET_RS08350 is actually a homoserine phosphotransferase. We cloned DET_RS08350 into a *thrB::kan* strain of E. coli and found that it rescued growth in minimal glucose medium with aTc: OD_600_ increased by 0.22-0.25 after 47 hours (3 replicates).

Second, we identified a potential phosphoserine aminotransferase in *Paucidesulfovibrio gracilis* (UniProt A0A1T4W7T3) that is 30% identical to the *serC3* from *C. botulinum* that we confirmed. We cloned the *P. gracilis* gene into a *serC::kan* strain of *E. coli* (Baba et al. 2006) and found that it rescued growth in minimal M9 medium with glucose and no aTc: OD_600_ increased by 0.25-0.29 after 48 hours (3 replicates). Similarly, expression of a diverged homolog from a metagenome-assembled genome of a *Methanomethylophilaceae sp.* (UniProt A0A843E9R6) rescued growth: after 72 hours in minimal M9 medium with glucose and aTc, OD_600_ increased by 0.14-0.19 (3 replicates).

## Appendix 6 Highly-diverged regulatory subunits of acetolactate synthase

We noticed that the regulatory subunit for acetolactate / acetohydroxybutanoate synthase appeared to be missing from many Thermoproteota, including from *Pyrolobus fumarii*, which is prototrophic for amino acids (Blöchl et al. 1997). In many Thermoproteota, we found an ACT domain protein conserved near the catalytic subunit. For instance, KCR_RS03285 from *Korarcheum cryptofilum* has an ACT domain and is encoded adjacent to the catalytic subunit (KCR_RS03290). When we used the AlphaFold 3 server (Abramson et al. 2024) to predict a protein complex between the two putative subunits from *K. cryptofilum*, it reported a high-confidence interaction (ipTM, the predicted template modeling score for the interface, was 0.83). Furthermore, the predicted interaction is similar to the experimentally determined interaction of the AHAS subunits from yeast (Figure S6). So, in the revised GapMind, KCR_RS03285 is included as a predicted regulatory subunit of AHAS.

**Figure S6:**
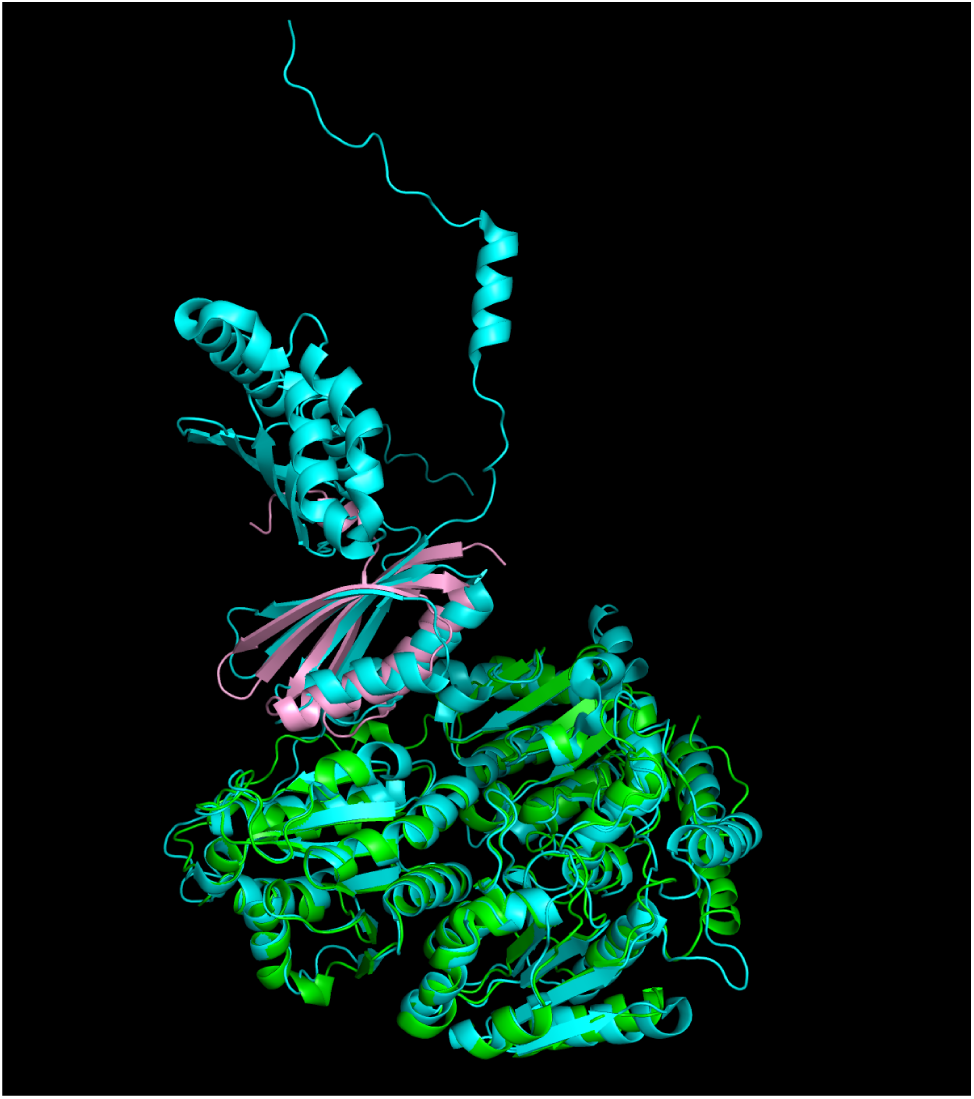
Acetolactate synthase (AHAS) from *Korarcheum cryptofilum* with a putative short regulatory subunit. The predicted structure for the *K. cryptofilum* enzyme (with catalytic subunit in green and regulatory subunit in pink) was aligned to the experimental structure for the yeast enzyme (in cyan) by using the align command of the PyMOL Molecular Graphics System (Version 2.5.2, Schrödinger, LLC). The yeast enzyme includes five copies of each subunit; only one copy of each is shown here (chains A and C from PDB:6U9D).

We identified another putative truncated regulatory subunit in many strains of *Chryseobacterium* (i.e., UniProt A0A316WEF5). This gene is encoded adjacent to the catalytic subunit, and is over 40% identical to the N-terminal part of a regulatory subunit from *Pedobacter* (CA265_RS15810, which was confirmed by fitness data). Because some of these strains of *Chryseobacterium* grow in the absence of any one of the branched-chain amino acids (i.e., GW821-FHT04C04 and GW821-FHT04A06; M. N. Price and H. P. Lesea, in preparation), the one-domain protein is presumably functional, and it is included in the revised GapMind as a prediction.

## Acknowledgements

We thank Hans Carlson for assisting with RB-TnSeq assays for *Rhodanobacter denitrificans*. This material by ENIGMA- Ecosystems and Networks Integrated with Genes and Molecular Assemblies (http://enigma.lbl.gov), a Science Focus Area Program at Lawrence Berkeley National Laboratory is based upon work supported by the U.S. Department of Energy, Office of Science, Office of Biological & Environmental Research under contract number DE-AC02-05CH11231.

